# Social Network Analysis of the Genealogy of Strawberry: Retracing the Wild Roots of Heirloom and Modern Cultivars

**DOI:** 10.1101/2020.09.30.320689

**Authors:** Dominique D.A. Pincot, Mirko Ledda, Mitchell J. Feldmann, Michael A. Hardigan, Thomas J. Poorten, Daniel E. Runcie, Christopher Heffelfinger, Stephen L. Dellaporta, Glenn S. Cole, Steven J. Knapp

**Affiliations:** Department of Plant Sciences, University of California, Davis, One Shields Avenue, Davis, California, 95616, USA; Department of Molecular, Cellular, and Developmental Biology, Yale University, New Haven, Connecticut, 06520, USA

**Keywords:** *Fragaria*, kinship, domestication, DNA forensics, biodiversity, conservation genetics

## Abstract

The widely recounted story of the origin of cultivated strawberry (*Fragaria* × *ananassa*) oversimplifies the complex interspecific hybrid ancestry of the highly admixed populations from which heirloom and modern cultivars have emerged. To develop deeper insights into the three century long domestication history of strawberry, we reconstructed the genealogy as deeply as possible—pedigree records were assembled for 8,851 individuals, including 2,656 cultivars developed since 1775. The parents of individuals with unverified or missing pedigree records were accurately identified by applying exclusion analysis to array-genotyped single nucleotide polymorphisms. We identified 187 wild octoploid and 1,171 *F.* × *ananassa* founders in the genealogy, from the earliest hybrids to modern cultivars. The pedigree networks for cultivated strawberry are exceedingly complex labyrinths of ancestral interconnections formed by diverse hybrid ancestry, directional selection, migration, admixture, bottlenecks, overlapping generations, and recurrent hybridization with common ancestors that have unequally contributed allelic diversity to heirloom and modern cultivars. Fifteen to 333 ancestors were predicted to have transmitted 90% of the alleles found in country-, region-, and continent-specific populations. Using parent-offspring edges in the global pedigree network, we found that selection cycle lengths over the last 200 years of breeding have been extraordinarily long (16.0-16.9 years/generation) but decreased to a present-day range of 6.0-10.0 years/generation. Our analyses uncovered conspicuous differences in the ancestry and structure of North American and European populations and shed light on forces that have shaped phenotypic diversity in *F.* × *ananassa*.

---

The strawberries found in markets around the world today are produced by cultivated strawberry (*Fragaria × ananassa* (Weston) Duchesne ex Rozier), a species domesticated over the last 300 years (Darrow 1966). *F. × ananassa* is technically not a species but an admixed population of interspecific hybrid lineages between cross-compatible wild allo-octoploid (2n = 8x = 56) species with shared evolutionary histories (Duchesne 1766; Darrow 1966; Liston *et al.* 2014). The earliest *F. × ananassa* cultivars originated as spontaneous hybrids between *F. chiloensis* and *F. virginiana* in Brittany, the Garden of Versailles, and other Western European gardens in the early 1700s, shortly after the migration of *F. chiloensis* from Chile to France in 1714 (Duchesne 1766; Bunyard 1917; Darrow 1966; Pitrat and Faury 2003). Their serendipitous origin was discovered by the French botanist Antoine Nicolas Duchesne (1747-1827) and famously described in a treatise on strawberries that biologists suspect included one of the first renditions of a phylogenetic tree (Duchesne 1766). Even though those studies pre-dated both the advent of genetics and the discovery of ploidy differences in the genus, the phylogenies were remarkably close to hypotheses that emerged more than 150 years later (Darrow 1966; Staudt 1988, 2003; Dillenberger *et al.* 2018). The early interspecific hybrids were observed to be phenotypically unique and horticulturally superior to their wild octoploid parents, which drove the domestication of *F. × ananassa*. Hardigan *et al.* (2020a,b) showed that hybrids between *F. chiloensis* and *F. virginiana* had nearly double the heterozygosity of their parents, which almost certainly boosted phenotypic variation and fueled *F. × ananassa* domestication. The cultivation of *F. × ananassa* steadily increased and ultimately supplanted the cultivation of other strawberry species, forever changing strawberry production and consumption worldwide (Fletcher 1917; Darrow 1966; Wilhelm and Sagen 1974; Finn *et al.* 2013).

The romanticized and widely recounted story of the origin of cultivated strawberry, while compelling, oversimplifies the complexity of the wild ancestry and 300-year history of domestication (Darrow 1966). The domestication of *F. × ananassa* has been documented in narrative histories and pedigree- and genome-informed studies of genetic diversity and population structure, but has not been fully untangled or deeply studied (Clausen 1915; Fletcher 1917; Darrow 1966; Wilhelm and Sagen 1974; Sjulin and Dale 1987; Bringhurst *et al.* 1990; Dale and Sjulin 1990; Johnson 1990; Sjulin 2006; Hancock *et al.* 2008; Horvath *et al.* 2011; Sánchez-Sevilla *et al.* 2015). The only pedigree-informed studies of the breeding history of cultivated strawberry focused on an analysis of the ancestry of 134 North American cultivars developed between 1960 and 1985 (Sjulin and Dale 1987; Dale and Sjulin 1990). They identified 53 founders in the pedigrees of those cultivars and estimated that 20 founders contributed approximately 85% of the allelic diversity. The inference reached in those studies and others was that cultivated strawberry is genetically narrow (Sjulin and Dale 1987; Dale and Sjulin 1990; Hancock and Luby 1995; Graham *et al.* 1996; Hancock *et al.* 2001; Hummer 2008; Gaston *et al.* 2020). The genetic narrowness hypothesis, however, has not been supported by genome-wide analyses of DNA variants, which have shown that *F. chiloensis*, *F. virginiana*, and *F. × ananassa* harbor massive nucleotide diversity and that a preponderance of the alleles transmitted by the wild octoploid founders have survived domestication and been preserved in the global *F. × ananassa* population (Hardigan *et al.* 2020a,b).

The domestication of cultivated strawberry has followed a path quite different from that of other horticulturally important species, many of which were domesticated over millennia and trace to early civilizations, e.g., apple (*Malus domestica*), olive (*Olea europaea* subsp. *europaea*), and wine grape (*Vitis vinifera* subsp. *vinifera*) (Purugganan and Fuller 2009; Myles *et al.* 2011; Meyer *et al.* 2012; Meyer and Purugganan 2013; Cornille *et al.* 2014; Larson *et al.* 2014; Diez *et al.* 2015; Duan *et al.* 2017). Although the octoploid progenitors were cultivated before the emergence of *F. × ananassa*, the full extent of their cultivation is unclear and neither appears to have been intensely domesticated, e.g., Hardigan *et al.* (2020b) did not observe changes in the genetic structure between land races and wild ecotypes of *F. chiloensis*, a species cultivated in Chile for at least 1,000 years (Finn *et al.* 2013). With less than 300 years of breeding, pedigrees for thousands of *F. × ananassa* individuals have been recorded, albeit in disparate sources. To delve more deeply into the domestication history of cultivated strawberry, we assembled pedigree records from hundreds of sources and reconstructed the genealogy as deeply as possible. One of the original impetuses for this study was to identify historically important and genetically prominent ancestors for whole-genome shotgun (WGS) resequencing and genome-scale analyses of nucleotide diversity (Hardigan *et al.* 2020a,b).

One challenge we faced when building the pedigree database and reconstructing the genealogy of strawberry was the absence of pedigree records for 96% of the 1,287 accessions preserved in the University of California, Davis (UCD) Strawberry Germplasm Collection, hereafter identified as the ‘California’ population. To solve this problem, authenticate pedigrees, and reconstruct the genealogy of the California population, we applied exclusion analysis in combination with high-density single nucleotide polymorphism (SNP) genotyping (Chakraborty *et al.* 1974; Elston 1986; Goldgar and Thompson 1988; Pena and Chakraborty 1994; Vandeputte 2012; Vandeputte and Haffray 2014). Here, we describe the accuracy of parent identification by exclusion analysis among individuals genotyped with 35K, 50K, or 850K SNP arrays (Bassil *et al.* 2015; Verma *et al.* 2016; Hardigan *et al.* 2020a). Several thousand SNP markers common to the three arrays were integrated to develop a SNP profile database for the exclusion analyses described here.

The genealogies (pedigree networks) of domesticated plants, especially those with long-lived individuals, overlapping generations, and extensive migration and admixture, can be challenging to visualize and comprehend (Mäkinen *et al.* 2005; Trager *et al.* 2007; Voorrips *et al.* 2012; Shaw *et al.* 2014; Fradgley *et al.* 2019; Muranty *et al.* 2020). We used Helium (Shaw *et al.* 2014) to visualize certain targeted pedigrees; however, the strawberry pedigree network was too large and complex to be effectively visualized and analyzed with traditional pedigree visualization approaches.

The pedigree networks of plants and animals share many of the features of social networks with nodes (individuals) connected to one another through edges (parent-offspring relationships) (Barabási *et al.* 2011; Barabási 2016; Contandriopoulos *et al.* 2018). We used social network analysis (SNA) methods, in combination with classic population genetic methods, to the analyze the genealogy and develop deeper insights into the domestication history of strawberry (Lacy 1989, 1995; Barabási *et al.* 2011; Barabási 2016; Contandriopoulos *et al.* 2018). SNA approaches have been applied in diverse fields of study but have apparently not yet been applied to the problem of analyzing and characterizing pedigree networks (Moreno 1953; Scott 1988; Edwards 1992; Wasserman and Faust 1994; Kominakis 2001). With SNA, narrative data (birth certificates and pedigree records) are translated into relational data (parent-offspring and other genetic relationships) and summary statistics (betweenness centrality and out-degree) and visualized as sociograms (pedigree networks) (Barabási *et al.* 2011; Barabási 2016; Contandriopoulos *et al.* 2018). Here, we report insights gained from studies of the formation and structure of domesticated populations world-wide, the complex wild ancestry of *F. × ananassa*, and genetic relationships among extinct and extant ancestors in demographically unique domesticated populations tracing to the earliest hybrids (Darrow 1966).

## Materials and Methods

### Pedigree Record Assembly, Documentation, and Annotation

We located and assembled pedigree records for strawberry accessions from more than 807 documents, databases, and other sources including: (a) US Patent and Trademark Office Plant Patents (https://www.uspto.gov/); (b) Germplasm Resource and Information Network (GRIN) passport data for accessions preserved in the USDA National Plant Germplasm System (NPGS; https://www.ars-grin.gov/); (c) the original unpublished UCD laboratory notebooks and other documents of Royce S. Bringhurst archived in a special collection at the Merill-Cazier Library, Utah State University, Logan, Utah (Bringhurst 1918-2016; USU COLL MSS 515; http://archiveswest.orbiscascade.org/ark:/80444/xv47241); (d) the original unpublished University of California, Berkeley (UCB) laboratory notebooks of Harold E. Thomas loaned by Phillip Stewart (Driscoll’s, Watsonville, California); (e) an obsolete electronic database discovered and recovered at UCD; (f) an electronic pedigree database for public cultivars developed by Thomas Sjulin, a former strawberry breeder at Driscoll’s, Watsonville, California; (g) scientific, technical bul-letins, and popular press articles; and (h) garden catalogs (Files S1-S3).

The pedigree records and other input data were manually curated and deduplicated. The database was constructed in a standard trio format (offspring, mother, father) with supporting passport data, which included: (a) alphanumeric identification numbers; (b) common names or aliases; (c) accession types (e.g., cultivars, breeding materials, or wild ecotypes); (d) birth years (years of origin); (e) geographic origin; (f) inventor (breeder or institution) names; (g) taxonomic classifications, and (h) DNA-authenticated pedigrees for genotyped UCD accessions, as de-scribed below (File S1). Because a parent could be a male in one cross and female in another, and parent sexes were frequently unknown or inconsistently recorded in pedigree records, the ‘mother’ (parent 1) and ‘father’ (parent 2) designations were arbitrary and unimportant to our study.

Germplasm accession numbers in the pedigree database included ‘plant introduction’ (PI) numbers for USDA accessions, UCD identification numbers for UCD accessions, and assorted other identification numbers. UCD accession numbers were written in a 10-digit machine-readable and searchable format to convey birth year and unique numbers, e.g., the UCD ID ‘65C065P001’ identifies a single individual (P001) in full-sib fam-ily C065 born in 1965 that was identified in historic records as ‘65.65-1’ (Bringhurst 1918-2016; Bringhurst *et al.* 1980). The latter is the ‘Bringhurst’ notation found in the historic pedi-gree records for UCD accessions and US Plant Patents. The decimals and dashes in the original notation created problems with data curation, analysis, and sorting. To solve this, the original ‘Bringhurst’ accession numbers (e.g., 65.65-1) were con-verted into the 10-digit machine-readable accession numbers (e.g., 65C065P001) reported in our pedigree database, where ‘C’ identifies a cultivated strawberry accession. Common names (aliases) of cultivars and accessions (if available) were concante-nated with underscores to create machine-readable and sortable names, e.g., the name for the *F. × ananassa* cultivar ‘Madame Moutot’ was stored as ‘Madame_Moutot’. Cultivars sharing names were made unique by appending an underscore and their year. Throughout the pedigree database, unknown individuals were created as necessary and identified with unique alphanu-meric identification numbers starting with the prefix ‘Unknown’, followed by an underscore, a species acronym when known or NA when unknown, an underscore, and consecutive numbers, e.g., ‘Unknown_FC_071’ identifies unknown *F. chiloensis* founder 71. The species acronyms applied in our database were FA for *F. × ananassa*, FC for *F. chiloensis*, FV for *F. virginiana*, FW for *F. vesca* (woodland strawberry), FI for *F. iinumae*, FN for *F. nipponica*, FG for *F. viridis* (green strawberry), FM for *F. moschata*, and FX for other wild species or interspecific hybrids, e.g., *F. × vescana*.

### Plant Material and SNP Profile Database

To develop a SNP profile database for DNA forensic and population genetic analyses (see below), we recalled and reanalyzed SNP marker genotypes for 1,495 individuals, including 1,235 UCD and 260 USDA accessions (asexually propagated individuals) previously genotyped by Hardigan *et al.* (2018) with the iStraw35 SNP array (Bassil *et al.* 2015; Verma *et al.* 2016). SNP marker genotypes were automatically called with the Affymetrix Axiom Analysis Suite (v1.1.1.66, Affymetrix, Santa Clara, CA). DNA samples with > 6% missing data were dropped from our analyses. We used quality metrics output by the Affymetrix Axiom Analysis Suite and custom R scripts and the R pack-age *SNPRelate* (Zheng *et al.* 2012) to identify and select codomi-nant SNP markers with genotypic clustering confidence scores (1 – *pC*) ≥ 0.01, where *p*_*C*_ is the posterior probability that the SNP genotype for an individual was assigned to the correct genotypic cluster (Affymetrix Inc. 2015). This yielded 14,650 high confidence co-dominant SNP markers for paternity-maternity analyses. While SNP markers are co-dominant by definition, a certain percentage of the SNP markers assayed in a popula-tion produce genotypic clusters lacking one of the homozygous genotypic clusters. These so-called ‘no minor homozygote’ SNP markers were excluded from our analyses.

For a second DNA forensic analysis, 1,561 UCD individuals were genotyped with 50K or 850K SNP arrays (Hardigan *et al.* 2020a). This study population included 560 hybrid offspring from crosses among 27 elite UCD parents, the *F. × ananassa* culti-var ‘Puget Reliance’, and the *F. chiloensis* subsp. *lucida* ecotypes ‘Del Norte’ and ‘Oso Flaco’. Hardigan *et al.* (2020a) included 16,554 SNP markers from the iStraw35 and iStraw90 SNP arrays on the 850K SNP array. To build a SNP profile database for the second paternity-maternity analysis, we identified 2,615 SNP markers that were common to the three arrays and produced well separated co-dominant genotypic clusters with high confidence scores (*pC* > 0.99) and < 6% missing data (Bassil *et al.* 2015; Verma *et al.* 2016; Hardigan *et al.* 2020a).

We subdivided the global population (entire pedigree) into ‘California’ and ‘cosmopolitan’ populations, in addition to continent-, region-, or country-specific populations, for different statistical analyses. These subdivisions are documented in the pedigree database (File S1). The California population included 100% of the UCD individuals (*n* = 3, 540) from the global popu-lation, in addition to 262 non-California individuals that were ascendants of UCD individuals. The cosmopolitan population included 100% of the non-California (non-UCD) individuals (*n* = 5, 193), in addition to 160 California individuals that were ascendants of non-California individuals. We subdivided individuals in the US population (excluding UCD individuals) into Midwestern, Northeastern, Southern, and Western US populations. The Western US population included only those UCD individuals that were ascendants in the pedigrees of Western US individuals. The country specific subdivisions were Australia, China, Japan, South Korea, Belgium, Czechoslovakia, Denmark, England, Finland, France, Germany, Israel, Italy, the Nether-lands, Norway, Poland, Russia, Scotland, Spain, Sweden, and Canada.

### DNA Forensic Analyses

We applied standard DNA forensic approaches for diploid organ-isms to the problem of identifying parents and authenticating pedigrees in allo-octoploid strawberry (Chakraborty *et al.* 1974; Elston 1986; Jones and Ardren 2003; Telfer *et al.* 2015; Muranty *et al.* 2020). Genotypic transgression ratios were estimated for all possible duos and trios of individuals in two study populations (described above) from genotypes of multiple SNP marker loci.

For duos of individuals in the SNP profile database for a population, the genotypic transgression score for the *i*th SNP marker was estimated by

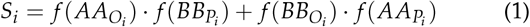

where *i* = 1, 2,…, *m*, *m* = number of SNP marker loci genotyped in each pair of probative DNA samples, *f* (– –*O*_*i*_) is the frequency of a homozygous genotype (coded *AA* and *BB*) in the candidate offspring individual and *f* (– –*P*_*i*_) is the frequency of a homozygous genotype in the candidate parent individual (similarly coded *AA* and *BB*) for the *i*th SNP marker locus. This equation was applied to a single pair of candidate individuals at a time and was thus constrained to equal 0 or 1; hence, *S*_*i*_ = 0 when homozygous genotypes were identical for a pair of individuals and *S*_*i*_ = 1 when homozygous genotypes were different for a pair of individuals. Duo-trangression ratios (*DTR*s) were estimated for every pair of individuals in the population by summing *S*_*i*_ estimates from equation (1) over *m* marker loci:

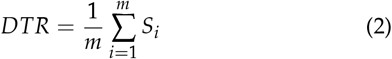

For trios of individuals in the SNP profile database for a population, the genotypic transgression score for the *i*th SNP marker was estimated by

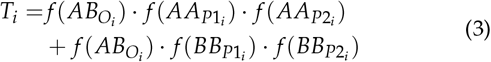

where 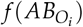 is the frequency of a heterozygous genotype (coded *AB*) in the candidate offspring individual, *f* (– –*P*1_*i*_) is the frequency of either homozygous genotype (*AA* or *BB*) in candidate parent 1 (*P*1), and *f* (– –*P*2_*i*_) is the frequency of either homozygous genotype in candidate parent 2 (*P*2) for the *i*th SNP marker locus. Trio transgression ratios (*TTR*s) were estimated for every trio of individuals in the population by summing *T_i_* estimates from equation (3) over *m* marker loci:

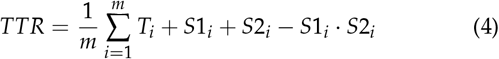

where *m* is the number of SNP marker loci genotyped for a trio of individuals, *S*1_*i*_ is the score estimated from equation (1) for candidate parent 1, and *S*2_*i*_ is the score estimated from equation (1) for candidate parent 2. To avoid double counting transgressions, *TTR* estimates were corrected by subtracting *S*1_*i*_ × *S*2_*i*_.

Our analyses yielded *DTR* and *TTR* estimates for paternity and maternity exclusion tests among genotyped individuals in the study populations. The putative parents of offspring were identified by estimating the probability of paternity (or maternity) from equations (2) and (4) and empirically estimating statistical significance thresholds by bootstrapping—50,000 bootstrap samples of size *n* were drawn with replacement from *n* probative DNA samples of individuals with declared parents in the population (Efron 1982; Simon and Bruce 1991; Manly 2006; Berry *et al.* 2014). The ‘declared’ or ‘stated’ parents are those recorded in pedigree records, whereas the ‘DNA-authenticated’ parents are those verified by exclusion analysis. The bootstrap-estimated *TTR*-threshold of *TTR* ≤ 0.01 yielded false-positive and negative probabilities of zero when estimated by summing *T_i_* estimates over 14,650 SNP marker loci. Similarly, the bootstrap-estimated *DTR*-threshold of *DTR* ≤ 0.0016 yielded a false positive probability of zero and a false negative probability of 5% when estimated by summing *S*_*i*_ estimates over 14,650 SNP marker loci.

### Social Network Analyses

The pedigree networks for global, California, and cosmopolitan populations were analyzed and visualized as directed social networks using the R package *igraph* (version 1.2.2; Csardi and Nepusz 2006), where every edge in the graph connects a parent node to an offspring node and information flows unidirectionally from parents to offspring (Wasserman and Faust 1994). The pedigree networks or sociograms were visualized using the open-source software Gephi (version 0.9.2; Bastian *et al.* 2009; https://gephi.org/). We estimated the number of edges (*d* = degree) and in-degree (*d_i_*), out-degree (*d_o_*), and betweenness centrality (*B*) statistics for every individual in a sociogram (Wasserman and Faust 1994). *d_i_* estimates the number of known parents, where *d_i_* = 0 when neither parent is known (for founders), 1 when one parent is known, and 2 when both parents are known. *d_o_* estimates the number of descendants of an individual. A ‘geodesic’ is the shortest path between two nodes in the network and estimates the number of generations in the pedigree of an individual (Hayes 2000). *D* is the longest geodesic in the network and estimates the largest number of generations for a descendant in the pedigree or the maximum depth of the pedigree (Hayes 2000). *B* estimates the connectivity of an individual to other individuals in a network (the number of geodesics connecting a node to other nodes), essentially the flow of information (alleles) and information ‘bottlenecks’ (Freeman 1977; Wasserman and Faust 1994; Yu *et al.* 2007; Pavlopoulos *et al.* 2011). *B* was estimated by

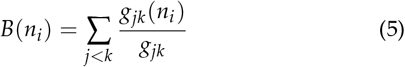

where *n*_*i*_ is the *i*th node (individual), *i*, *j*, and *k* are different nodes, *g*_*jk*_ is the number of geodesics occurring between nodes *j* and *k*, and *g*_*jk*_ (*n*_*i*_) is the number of geodesics that pass through the *i*th node (Freeman 1977; Wasserman and Faust 1994; Brandes 2001; Csardi and Nepusz 2006). *B* = 0 when *d*_*i*_ or *d*_*o*_ equal zero.

Standard social network analysis metrics and terminology were used to classify individuals and describe their importance in the genealogy, which are analogous to applications in diverse fields of study (Gursoy *et al.* 2008; Koschützki and Schreiber 2008; Morselli 2010; Kim and Song 2013; Nerghes *et al.* 2015). Using *B* and *d_o_* estimates, ancestors were classified as globally central 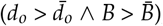, locally central 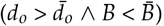, broker 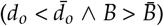, or marginal 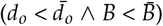.

### Selection Cycle Length Calculations

The pedigree network for every cultivar was extracted from the global pedigree network and included the cultivar (the youngest terminal node) and every ascendant (founder and non-founder) of the cultivar. Selection cycle lengths (*S* = years/generation) were estimated for every cultivar by tracing every possible path (back in time) in the pedigree network from the cultivar to founders and calculating birth year differences for every parent-offspring edge (*y*_*i*_) in the path, where *y*_*i*_ is the number of years separating the *i*th parent-offspring edge. The mean selection cycle length was estimated by 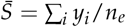, where *y*_*i*_ is the birth year difference for the *i*th parent-offspring edge, *n_e_* is the number of parent-offspring edges and *i* = 1, 2,…, *n*_*e*_. To understand how selection cycle length changed over time, we considered all 14,275 unique parent-offspring edges available in the pedigree, among which 9,486 had birth years known for both the parent and the offspring. For each edge, we computed its midpoint as the average birth year between the parent and the offspring and its size, i.e. the selection cycle length (*S*), as the difference in birth years between the parent and the offspring.

### Estimation of Coancestry and Pedigree-Genomic Relationship Matrices

The kinship or coancestry matrix (*A*) was estimated for the entire pedigree (*n* = 8, 851 individuals) using the *create.pedigree* and *kin* functions in the R package *synbreed* (version 0.12-12; Wimmer *et al.* 2012), where the *i*th diagonal element of *A* is the coefficient of coancestry of individual *i* with itself (*C*_*ii*_) and the *ij*th off-diagonal element of *A* is the coefficient of coancestry between individuals *i* and *j* (*C*_*ij*_) (Lynch and Walsh 1998). The genomic relationship matrix (*G*) was estimated for 1,495 individuals genotyped with 14, 650 SNP markers selected to have minor allele frequencies (MAF) ≥ 0.05 and ≤ 10% missing data. *G* was estimated as described by VanRaden (2008) using the function *A.mat* in the R package *rrBLUP* (version 4.6.1; Endelman 2011). Missing genotypes were imputed using the mean genotype for each SNP marker.

We estimated the combined pedigree-genomic relationship matrix (*H*) for the entire pedigree (*n* = 8, 851 individuals) as described by Legarra *et al.* (2009). The *A* matrix was partitioned into four sub-matrices (*A*_11_, *A*_12_, *A*_21_, *andA*_22_), where the subscript 1 indexes ungenotyped and 2 indexes genotyped individ-uals. *G* and *A*_22_ had the same dimensions but different scales. To construct the scaled *G* matrix (Christensen 2012; Christensen *et al.* 2012; Gao *et al.* 2012), the mean of off-diagonal elements of 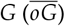 were scaled to match 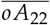 and the mean of diagonal elements of 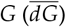 were scaled to match 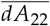:

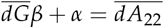

and

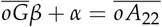

with scalar solutions

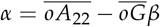

and

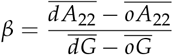

The *H* matrix was estimated using the scaled *G* matrix 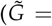 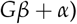 as described by Legarra *et al.* (2009):

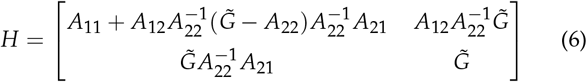

The open-source R code we developed to estimate *H* has been deposited in a FigShare database (File S6).

To study genetic relationships among extinct and extant individuals, we estimated separate *H* matrices for the California and cosmopolitan populations and applied principal component analysis (PCA) to the unscaled *H* matrices. Principal components were estimated by spectral decomposition of *H* using the *eigen* function from base R (version 4.0.0), which yielded eigenvalues, eigenvectors, and component scores. Scores for the first two principal components were then plotted using the R package *ggplot2* (Wickham 2016).

### Genetic Contributions of Founders and Ancestors

Coancestry or kinship (*A*) matrices were estimated for indi-viduals within continent-, region-, and country-specific focal populations using the *create.pedigree* and *kin* functions in the R package *synbreed* (version 0.12-9; Wimmer *et al.* 2012). Focal populations consisted of cultivars and their ascendants (ancestors). Founders are ancestors with unknown parents, which were assumed to be unrelated (Lacy 1989, 1995; Hartl and Clark 2007), whereas non-founders are ancestors with known parents. Terminal nodes in a pedigree network (sociogram) are either founders or the youngest descendants. The mean kinskip between the *i*th founder and cultivars in a focal population was estimated by

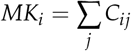

where *C*_*ij*_ = the kinship coefficient between the *i*th founder and *j*th cultivar in a focal population, *i* = 1, 2, ․ ․ ․, *n*, *j* = 1, 2, ․ ․ ․, *k*, *n* = the number of founders in the focal population, and *k* = the number of cultivars in the focal population. (Lacy 1989, 1995; Lynch and Walsh 1998; Hartl and Clark 2007). The proportional genetic contribution of the *i*th founder to a focal population was estimated by *P*_*i*_ = *MK*_*i*_ / ∑_*i*_ *MK*_*i*_. The number of founder equivalents (*Fe*) was estimated by *F*_*e*_ = 1/∑_*i*_ *MK*_*i*_, where *i* ∈ {founder_1_, founder_2_, ․․․, founder_*n*_} (Lacy 1989, 1995). Founder equivalents “are the number of equally contributing founders that would be expected to produce the same genetic diversity as in the population under study” (Lacy 1989).

The genetic contributions (*GC*) of ancestors (founders and non-founders) to a focal population were estimated by constructing a directed distance matrix (*D*) with dimensions identical to *A* (*n × n*) such that parents appeared in the matrix before offspring (alleles flow from parents to offspring but not *vice versa*). We used the directed distance (the number of parent-offspring edges between two accessions) to modify *A* so that coancestry coefficients were only estimated between ancestors and direct path cultivars. The directed distance matrix *D* was estimated using the *distances* function in the R package *igraph* (version 1.2.5; Csardi and Nepusz 2006), where non-zero distances in the *D* matrix were set equal to one. Coancestry coefficients for ascendants with no direct path to a cultivar were set equal to zero by taking the Hadamard product to generate the corrected coancestry matrix *A*^*^= *A* ⊙ *D*, where element *C*_*ii*_ = the coancestry coefficient for individual *i* with itself (Hartl and Clark 2007). To estimate *GC* for each ancestor, we applied an iterative approach that entailed: (i) computing *D*, *A*, and *A*^*^ = *A* ⊙ *D* from the current pedigree; (ii) estimating *MK*_*i*_ for each ancestor; (iii) ranking *MK*_*i*_ estimates from largest to smallest; (iv) setting *GC*_*i*_ = *MK*_*i*_ for the ancestor with the largest *MK*_*i*_ estimate; (v) deleting the ancestor with the largest *MK*_*i*_ estimate and rebuilding the pedigree; and (vi) repeating the previous steps until genetic contributions (*GC*_*i*_) had been estimated for each ancestor. The proportional genetic contribution of the *i*th ancestor to a focal population was estimated by *P*_*i*_ = *GC*_*i*_ / ∑_*i*_ *GC*_*i*_.

### Data Availability

File S1 contains the pedigree database with parents and offspring in a standard trio format (offspring, mother, father) with the following passport data: (a) alphanumeric identification number; (b) common names or aliases; (c) accession types (e.g., cultivars, breeding materials, or wild ecotypes); (d) birth years (years of origin); (e) geographic origins; (f) inventor (breeder or institution) names; (g) taxonomic classifications, and (h) DNA-authenticated pedigrees for genotyped UCD accessions. File S2 contains pedigrees of in the Helium format with parents and offspring identified by common names or aliases (Shaw *et al.* 2014; https://github.com/cardinalb/helium-docs/wiki). File S3 is a complete bibliography of the databases and documents we referenced to build the pedigree database. Files S4 and S5 contain betweenness (*B*), in-degree (*d*_*i*_), and out-degree (*d*_*o*_) statistics, structural role assignments, giant or halo component assignments, and coancestry-based estimates of the genetic contributions of founders and ancestors to cultivars in the California and Cosmopolitan populations, respectively. File S6 contains R code developed to estimate *H* from *A* and *G* as described by Legarra *et al.* (2009). The example input files from Legarra *et al.* (2009) for computing the *H* matrix are included. File S7 contains R code developed for exclusion (paternity-maternity) analyses. Table S1 details the most prominent ecotype founders and their coancestry-based estimates of genetic contribution to the California and Cosmopolitan populations. All supplements were uploaded to the FigShare Data Repository.

## Results and Discussion

### Genealogy of Cultivated Strawberry

We reconstructed the genealogy of cultivated strawberry as deeply as possible from wild founders to modern cultivars (Fig. 1; File S1). To build the database, pedigree records for 8,851 individuals were assembled from more than 800 documents including scientific and popular press articles, laboratory note-books, garden catalogs, cultivar releases, plant patent databases, and germplasm repository databases (Fig. 1; see File S3 for a complete bibliography). The database holds pedigree records and passport data for 2,656 *F. × ananassa* cultivars, of which approximately 310 were private sector cultivars with pedigree records in public databases (File S1). The parents of the private sector cultivars, however, were nearly always identified by cryptic alphanumeric codes, and thus could not be integrated into the ‘giant component’ of the sociogram (pedigree network) (Fig. 1).

**Figure 1.**
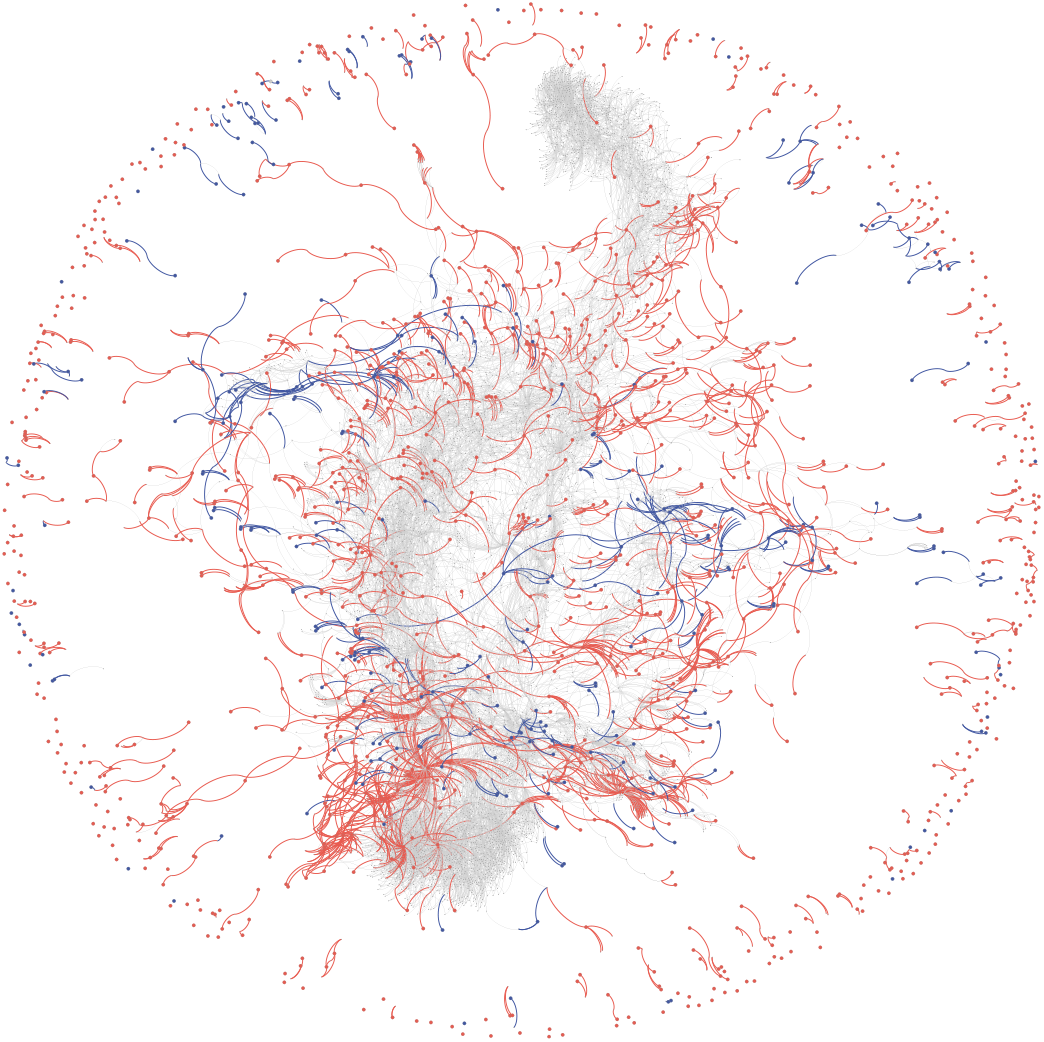
Global Pedigree Network for Cultivated Strawberry. So-ciogram depicting ancestral interconnections among 8,851 ac-cessions, including 8,424 *F. × ananassa* individuals originating as early as 1775, of which 2,656 are cultivars. The genealogy includes *F. chiloensis* and *F. virginiana* founders tracing to 1624 or later. Nodes and edges for 267 wild species founders are shown in blue, whereas nodes and edges for 1,171 *F. × ananassa* founders are shown in red. Founders are individuals with un-known parents. Nodes and edges for descendants (non-founders) are shown in light grey. The outer ring (halo of nodes and edges) are orphans or individuals in short dead-end pedigrees disconnected from the principal pedigree network or so-called ‘giant component’.

The global population was subdivided into ‘cosmopolitan’ and ‘California’ populations to delve more deeply into their unique breeding histories (Hardigan *et al.* 2020b; Fig. 1-2). This split was informed by demography and geography, insights gained from genome-wide analyses of nucleotide diversity and population structure (Hardigan *et al.* 2020a,b), and earlier DNA marker-informed studies of genetic diversity (Horvath *et al.* 2011; Sánchez-Sevilla *et al.* 2015; Hardigan *et al.* 2018). The cosmopolitan population included 100% of the non-California (non-UCD) individuals (*n* = 5, 193) from the global population, in addition to 160 California individuals identified as ascendants of non-California individuals. The non-California cultivar ‘Cascade’ (PI551759), for example, is a descendant of a cross between the California cultivar ‘Shasta’ (PI551663) and non-California cultivar ‘Northwest’ (PI551499) (https://www.ars.usda.gov/); hence, ‘Shasta’ was included in both the cosmopolitan and California populations. Similarly, the California population included 100% of the UCD individuals (*n* = 3, 540) from the global population, in addition to 262 non-California individuals that were identified as ascendants of UCD individuals. We nearly completely reconstructed the genealogy of the California population; however, as described below, pedigree records were missing for nearly every individual in the California population but were accurately ascertained using computer and DNA forensic approaches.

### Social Network Analyses Uncover Distinctive Differences in the Domestication History of California and Cosmopolitan Populations

We estimated that 80-90% of the individuals in the California and cosmopolitan pedigree networks were extinct (Fig. 2). Using SNP-array genotyped individuals preserved in public germplasm collections as anchor points, we searched for evidence that the allelic diversity transmitted by extinct founders had been ‘lost’. This is a difficult question to answer with certainty; however, the findings reported here, combined with the findings of Hardigan *et al.* (2020b), suggest that genetic diversity has been exceptionally well preserved in domesticated populations. Using SNA and principal component analyses (PCAs) of *H*, we did not observe structural features in sociograms or PCA plots that were indicative of the loss of novel ancestral genetic diversity (Fig 2). The kinship or numerator relationship matrix (*A*) was estimated for the entire pedigree of genotyped and ungenotyped individuals (VanRaden 2008; Legarra *et al.* 2009). For the present study, 1,495 historically important and geographically diverse UCD and USDA *F. × ananassa* individuals were genotyped with high-density SNP arrays (Bassil *et al.* 2015; Verma *et al.* 2016; Hardigan *et al.* 2020a). The genomic relationship matrix (*G*) was estimated for the genotyped individuals and combined with the *A* matrix to estimate the *H* matrix for the entire pedigree (Legarra *et al.* 2009). The global *H* matrix was partitioned as needed for subsequent analyses (Fig. 2).

**Figure 2.**
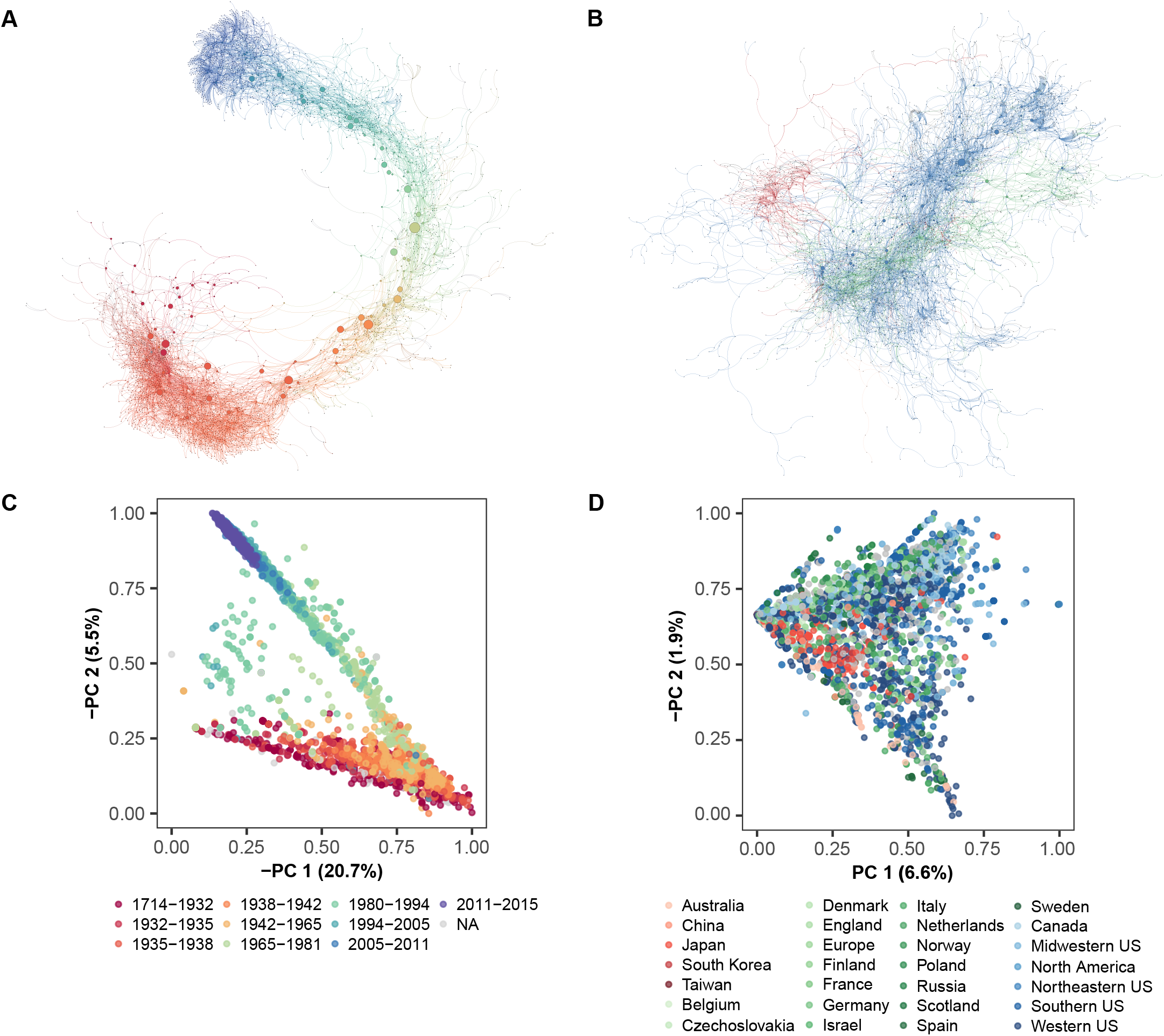
Genealogy for California and Cosmopolitan Populations of Cultivated Strawberry. (A) Sociogram depicting ancestral interconnections among 3,802 individuals in the ‘California’ population. This population included 3,452 *F. × ananassa* individuals developed at the University of California, Davis (UCD) from 1924 to 2012, in addition to 151 non-UCD *F. × ananassa* ascendants that originated between 1775 and 1924. Node and edge colors depict the year of origin of the individual in the pedigree network from oldest (red) to youngest (blue) with a continuous progression from warm to cool colors as a function of time (year of origin). Nodes and edges for individuals with unknown years of origin are shown in grey. (B) Sociogram depicting ancestral interconnections among 5,354 individuals in the ‘cosmopolitan’ population. This population included 5,106 *F. × ananassa* individuals developed across the globe between 1775 and 2018 and excludes UCD individuals other than UCD ancestors in the pedigrees of non-UCD individuals. Node and edge colors depict the continent where individuals in the pedigree network originated: Australia (orange), Asia (red), North America (blue), and Europe (green). Nodes and edges for individuals of unknown origin are shown in grey. (A and B) For both sociograms, node diameters are proportional to the betweenness centrality (*B*) metrics for individuals (nodes). Orphans and short dead-end pedigrees that were disconnected from the principal pedigree network (’giant component’) are not shown. (C) Principal component analysis (PCA) of the pedigree-genomic relationship matrix (*H*) for the California population. The *H* matrix (8,851 × 8,851) was estimated from the coancestry matrix (*A*) for 8,851 individuals and the genomic relationship matrix (*G*) for 1,495 individuals genotyped with a 35K SNP array. The PCA plot shows PC1 and PC2 coordinates for 3,802 individuals in the California population color coded by year-of-origin. (D) PCA of the *H* matrix for the cosmopolitan population. The PCA plot shows PC1 and PC2 coordinates for 5,354 individuals in the cosmopolitan population color coded by country, region, or continent of origin.

PCAs of the *H* matrices yielded two-dimensional visualizations of genetic relationships that were remarkably similar in shape and structure to sociograms for the California and cosmopolitan populations (Fig 2). We observed distinctive differences in the shapes and structures of the sociograms and PCA plots between the populations (Fig 2). The pattern in the cosmopolitan population was characteristic of pervasive admixture among individuals across geographies (Fig 2B and D). We observed a strong chronological trend in the California population (Fig 2A and C) but not in cosmopolitan population (Fig 2B and D). We observed a mid-twentieth century bottleneck in the California population (the sharp interior angle in the V-shaped structure of the PCA plot), in addition to a bottleneck pinpointed to approximately 1987-1993 when the California population became closed. We discovered that 48 founders contributed 100% of the allelic diversity to the California population from 1987 onward (Fig 2A and C; S1 File). Hardigan *et al.* (2020b) showed that even though nucleotide diversity had been progressively reduced by bottlenecks and selection, significant nucleotide diversity has persisted in the California population but was found to be unevenly distributed across the genome.

### DNA Forensic Approaches for Parent Identification and Pedigree Authentication in Octoploid Strawberry

When this study was initiated in early 2015, 1,235 *F. × ananassa* germplasm accessions (asexually propagated individuals) were preserved in the UCD Strawberry Germplasm Collection. The collection included 68 UCD cultivars with known pedigrees; however, pedigree records for the other 1,184 UCD individ-uals were unavailable. Using computer forensic approaches, pedigree records for 1,002 individuals were recovered from an obsolete electronic database. Because the authenticity and accuracy of those records were uncertain, every individual was genotyped with the iStraw35 SNP array to build a SNP profile database for parent identification by exclusion analysis (Jones and Ardren 2003; Vandeputte 2012; Vandeputte and Haffray 2014; Bassil *et al.* 2015; Verma *et al.* 2016). SNP marker genotypes were automatically called using the Affymetrix Axiom Suite, then manually curated to identify and extract codominant SNP markers with well separated genotypic clusters. This yielded 14,650 SNP markers for exclusion analyses. Genotyping errors were negligible (0.06-0.37%) and genotype-matching percentages for array-genotyped SNPs ranged from 99.63 to 99.95% among biological and technical replicates.

We estimated duo transgression ratios (*DTRs*) for all possi-ble paris or duos (761, 995) of individuals (Fig. 3). Trio transgression ratios (*TTRs*) were estimated for all possible triplets or trios of individuals with *DTR* estimates in the 0.00 to 0.01 range—individuals with *DTR* estimates > 0.0016 were excluded as parents (Fig. 3). For trio analyses, we included the possibility that offspring could arise by self-pollination, which yielded *n* × (*n* – 1) = 1,235 × 1,234 = 1,523,990 possible trios. Although this possibility does not arise in human or animal parent identification problems (Jones and Ardren 2003; Vandeputte 2012), offspring can arise from self-pollination in cultivated strawberry and other self-compatible plant species. The number of possible trios arising from crosses between two parents in the reference population was (*n* × [*n* – 1]) + (*n* × [*n* – 1] × [*n* – 2])/2 = 941,063,825.

**Figure 3.**
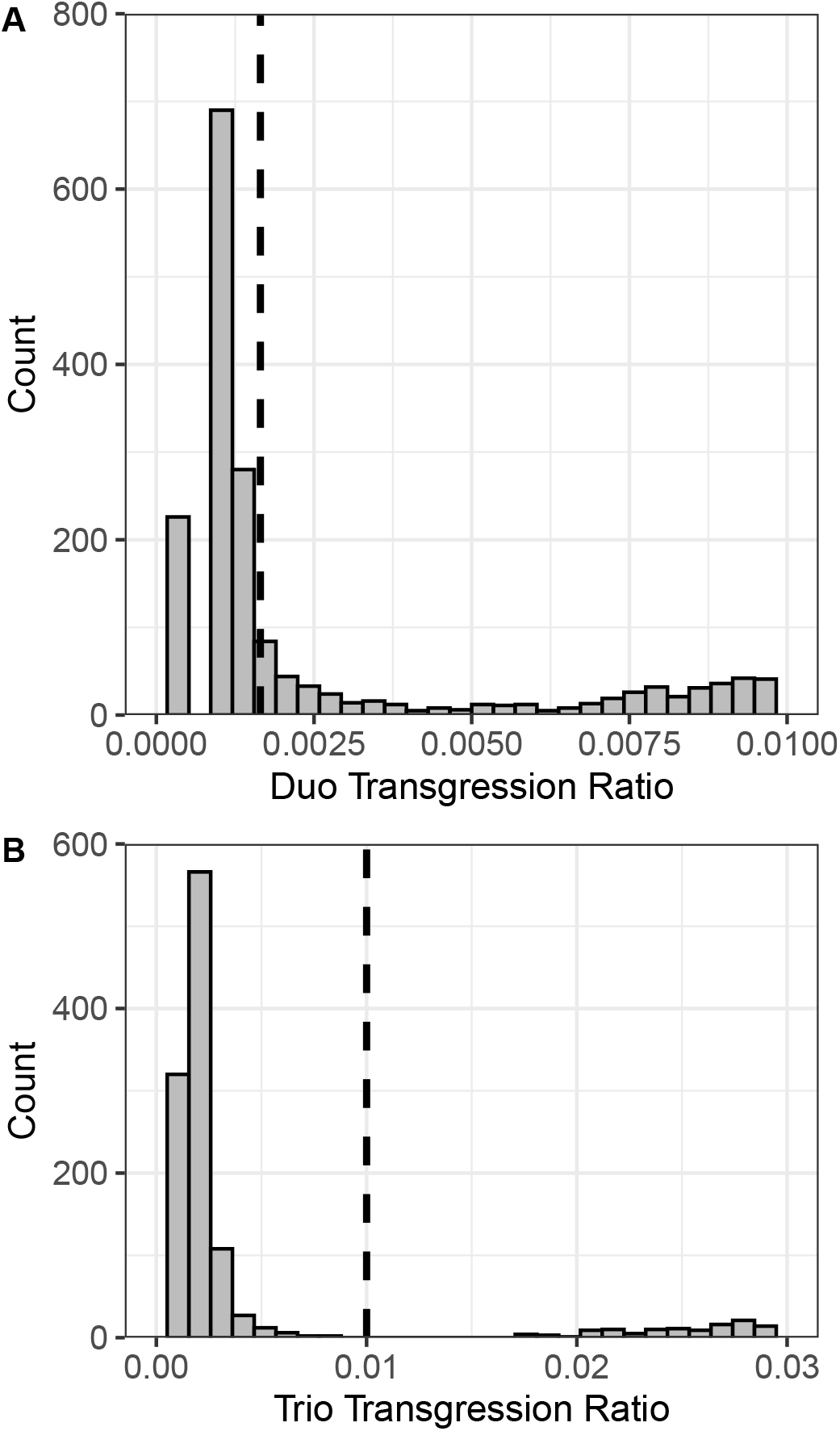
Exclusion Analyses. (A) Distribution of 2,708 duo transgression ratio (*DTR*) estimates falling in the 0.0 to 0.01 range. There were 761,995 possible duos among 1,235 individuals in the California population (*DTR* estimates > 0.01 are not shown). The vertical dashed line demarcates the bootstrap-estimated significance threshold (*DTR* < 0.0016) chosen to minimize false positives and negatives. (B) Distribution of 2,815 trio transgression ratio (*TTR*) estimates falling in the 0.00 to 0.03 range. There were 941,063,825 possible *TTR* estimates for trios among 1,235 individuals in the California population, which included 1,235 × 1,234 = 1,523,990 possible trios for offspring arising from self-pollination (*TTR* estimates > 0.03 are not shown). The vertical dashed line demarcates the bootstrap-estimated significance threshold (*TTR* < 0.01) chosen to minimize false positives and negatives. (A) and (B) *DTR*s and *TTR*s were estimated by summing over 14,650 SNP markers. Statistical significant thresholds for parent inclusion were empirically estimated from 50,000 boot-strap samples.

Trio exclusion analysis identified the parents of 1,044 UCD individuals with 100% accuracy and zero false positives–SNP profiles for both parents were present in the database for these individuals (Fig. 3). *DTR* estimates for parents with statistically significant *TTR* estimates (*TTR* < 0.01) were statistically significant (*DTR* < 0.0016). When the SNP profile for only one parent was present in the database (134 out of 1,235 individuals), duo exclusion analysis identified those parents with 95% accuracy and zero false positives (Fig. 3). When the DNA profile for only one parent exists in the database, the probability of a false negative slightly increases and the power to unequivocally identify that parent slightly decreases (Vandeputte 2012; Vandeputte and Haffray 2014). The difference in statistical power between the duo and trio method stems from differences in statistical power that arise from the presence of SNP profiles for both parents (*TTR*) as opposed to one parent (*DTR*) in the reference database (Elston 1986; Goldgar and Thompson 1988). For a diploid (or allo-polyploid) organism genotyped with bi-allelic subgenome-specific DNA markers, two out of nine possible genotypic combinations are informative for duo exclusion analysis, whereas 12 out of 27 possible genotypic combinations are informative for trio exclusion analysis (Vandeputte 2012; Vandeputte and Haffray 2014). Moreover, trio exclusion analysis includes two highly informative (statistically powerful) combinations where the candidate offspring are heterozygous (*AB*) and both parents are homozygous for the same allele (either *AA* or *BB*).

Our computer forensic search did not recover pedigree records for 220 individuals in the UCD population; however, we suspected that their parents might be present in the SNP profile database. Using duo and trio exclusion analyses, we identified both parents for 214 individuals and one parent each for the other six individuals. Hence, using a combination of computer and DNA forensic approaches, 2,222 out of 2,470 possible parents of 1,235 individuals (90.0%) in the UCD population were identified and documented in the pedigree database (File S1; Fig. 2). The parents declared in pedigree records (if known), identified by DNA forensic methods (if conclusive), or both are documented in the pedigree database (File S1). Despite their historic and economic importance, the pedigrees of individuals preserved in the UCD Strawberry Germplasm Collection had not been previously documented. Besides reconstructing the genealogy of the UCD population, previously hidden or unknown pedigrees of extinct and extant individuals were discovered in the historic UCD records of Harold E. Thomas, Royce S. Bringhurst, and others (Bringhurst 1918-2016; Bringhurst *et al.* 1990; Johnson 1990) and integrated into the global pedigree database (File S1).

To further validate the accuracy of DNA forensic approaches for parent identification in octoploid strawberry, we applied exclusion analysis to a population of 560 hybrid individuals developed from crosses among 30 UCD individuals (parents). The parents and hybrids and 1,561 additional UCD individuals were genotyped with 50K or 850K SNP arrays (Hardigan *et al.* 2020a). The 50K array was developed with SNP markers from the 850K array (Hardigan *et al.* 2020a), which included a subset of 16,554 legacy SNP markers from the iStraw35 and iStraw90 arrays (Bassil *et al.* 2015; Verma *et al.* 2016). We developed an integrated SNP profile database using 2,615 SNP markers common to the three arrays. Using parent-offspring trios, we discovered that the SNP profile for one of the parents (11C151P008) was a mismatch, whereas the SNP profiles of the other 29 parents perfectly matched their pedigree (birth) records. We discovered that the parent stated on the birth certificate for 11C151P008 was correct, but that the DNA sample and associated SNP marker profile were incorrect. Hence, the DNA sample mismatch was traced by exclusion analysis to a single easily corrected laboratory error.

These results highlight the power and accuracy of diploid Mendelian exclusion analysis for pedigree authentication (paternity and maternity analysis), intellectual property protection, and quality control monitoring of germplasm and nursery stock collections in octoploid strawberry using subgenome-specifc DNA markers. The application of these approaches was straight-forward because of the simplicity and accuracy of paralog- or homeolog-specific genotyping approaches in octoploid strawberry populations (Hardigan *et al.* 2020a). The development and robustness of subgenome-specific genotyping approaches has enabled the application of standard diploid genetic theory and methods in octoploid strawberry, including the exclusion analysis methods applied in the present study (Jones and Ardren 2003; Vandeputte 2012; Vandeputte and Haffray 2014; Fig. 3). The power and accuracy of these methods were rigorously tested and affirmed in a court of law where DNA forensic evidence was pivotal in proving the theft of University of California intellectual property (strawberry germplasm) by the defendants in a 2017 case in US District Court for the Northern District of California captioned *The Regents of the University of California v California Berry Cultivars, LLC, Shaw, and Larson* (Chivvis 2017). The DNA forensic approach and evidence in that case are documented in a publicly available expert report identified by case number 3:16-cv-02477 (https://ecf.cand.uscourts.gov/cgi-bin/login.pl).

### The Wild Roots of Cultivated Strawberry

Our genealogy search did not uncover pedigree records for *F. × ananassa* cultivars developed between 1714 and 1775, the 61 year period following the initial migration of *F. chiloensis* ecotypes from Chile to Europe (Duchesne 1766; Darrow 1966). The scarcity of pedigree records from the eighteenth century was anticipated because the interspecific hybrid origin of *F. × ananassa* was not discovered until mid-1700s (Duchesne 1766). ‘Madame Moutot’ was the only cultivar in the database with ancestry that could be directly traced to one of the putative original wild octoploid progenitors of the earliest *F. × ananassa* hybrids that emerged in France in the early 1700s (Fig. 4). Although the genealogy primarily covers the last 200 years of domestication and breeding (File S1), ascendants in the pedigree of the culti-var ‘Madame Moutot’ (circa 1906) traced to ‘Chili de Plougastel’ (Fig. 4), a putative clone of one of the original *F. chiloensis* subsp. *chiloensis* plants imported from Chile to France by the explorer Amédée-François Frézier (Gloede 1865; Carriére 1879; Bunyard 1917; Darrow 1966; Pitrat and Faury 2003). These plants were carried aboard the French frigate ‘St. Joseph’, delivered by Frézier to Brest, France (Bunyard 1917), and shared with Antoine Laurent de Jussieu, a botanist at the Jardin des plantes de Paris. According to de Lambertye (1864), the Frézier clone was widely disseminated and cultivated in Plougastel near Brest and inter-planted with *F. virginiana* (Duchesne 1766; Bunyard 1917; Pitrat and Faury 2003). Hence, some of the earliest spontaneous hybrids between *F. chiloensis* and *F. virginiana* undoubtedly arose in the strawberry fields of Brittany in the early 1700s (de Lambertye 1864; Darrow 1966; Pitrat and Faury 2003). The French naturalist Bernard de Jussieu, the brother of Antoine Laurent de Jussieu and a mentor of Antoine Duchesne—“the father of the modern strawberry”—brought clones of the original Frézier *F. chiloensis* plants to the Jardins du Château de Versailles (Gardens of Versailles) where Duchesne (1766) unraveled the interspecific hybrid origin of *F. × ananassa* (Darrow 1966; Williams 2001). The next earliest *F. chiloensis* founders appear to be a California eco-type identified in German breeding records from the mid-1800s and an anonymous ecotype in the pedigree of the French cultivar ‘La Constante’ from 1855 (Files S1-S2; Gloede 1865; Merrick 1870; Darrow 1937, 1966; Wilhelm and Sagen 1974).

**Figure 4.**
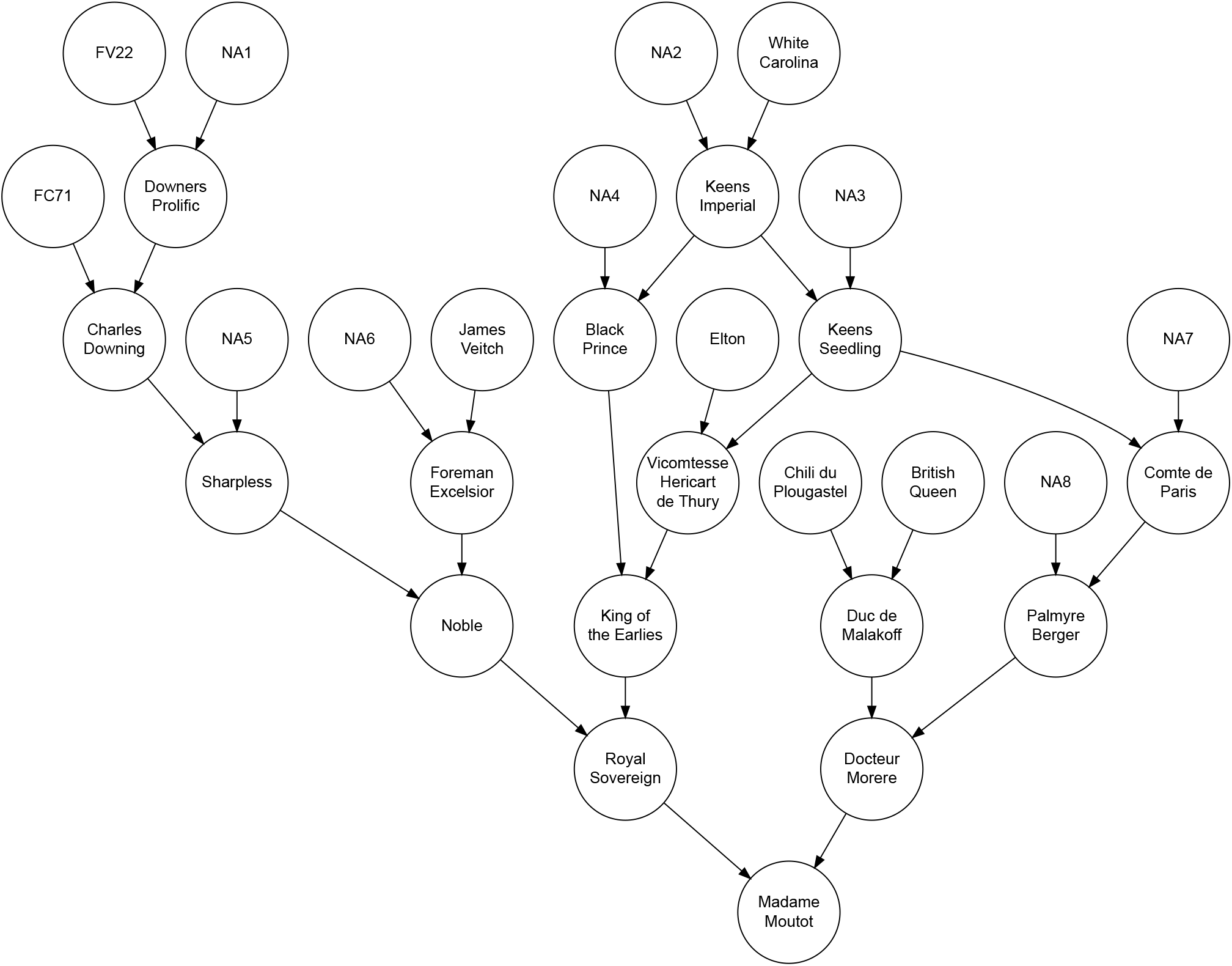
Pedigree for the Heirloom Cultivar ‘Madame Moutot’ (circa 1906). Arrows indicate the flow of genes from parents to offspring. FV22 is an unknown *F. virginiana* ecotype, FC71 is an unknown *F. chiloensis* ecotype, and ‘Chili du Plougastel’ is purportedly one of the original *F. chiloensis* individuals imported by Amédée-François Frézier from Chile to France in 1714. Unknown parents of individuals in the pedigree are identified by NA1, NA2,…, NA7. Terminal individuals in the pedigree are founders (individuals with unknown parents). The oldest *F. × ananassa* cultivar in the pedigree is ‘White Carolina’ (PI551681), which originated sometime before 1775.

The origins and identities of the earliest *F. virginiana* founders of *F. × ananassa* remain a mystery because their migrations from North America to Europe in the early 1600s and subsequent intra-continental migrations were not well documented (File S1; Duchesne 1766; de Lambertye 1864; Darrow 1937). The oldest *F. virginiana* individuals identified in historic documents and pedigree records were ‘Large Early Scarlet’ (1624), ‘Old Scarlet’ (1625), and ‘Hudson Bay’ (1780), all extinct (File S1). We identi-fied 30 anonymous *F. virginiana* and 76 anonymous *F. chiloensis* founders in the pedigree records. These individuals were as-signed unique alphanumerical aliases to facilitate reconstruction of the genealogy, e.g., FV22 is the alias for an anonymous *F. virginiana* founder and FC71 is the alias for an anonymous *F. chiloensis* founder in the pedigree of ‘Madame Moutot’ (Fig. 4; File S1).

### The Complex Hybrid Ancestry of Cultivated Strawberry

Once the interspecific hybrid origin of *F. × ananassa* became widely known (Duchesne 1766), domestication began in earnest with extensive intra-and interspecific hybridization, artificial se-lection, and intra-and intercontinental migration (Merrick 1870; Fletcher 1917; Darrow 1937). These forces shaped the genetic structure of the *F. × ananassa* populations that emerged in Europe and North America and ultimately migrated around the globe (Fletcher 1917; Darrow 1966; Sjulin and Dale 1987; Johnson 1990; Sjulin 2006; Horvath *et al.* 2011; Sánchez-Sevilla *et al.* 2015; Hardigan *et al.* 2018, 2020b). Over the next 250 years, horticulturalists and plant breeders repeatedly tapped into the wild reservoir of genetic diversity, especially wild octoploid taxa native to North America (Fig. 1; Table 1). There are numerous narrative accounts of what transpired, especially in Europe, North America, and California (Clausen 1915; Darrow 1937, 1966; Sjulin and Dale 1987; Bringhurst *et al.* 1990; Dale and Sjulin 1990; Johnson 1990; Hancock *et al.* 2001; Sjulin 2006; Hancock *et al.* 2010; Horvath *et al.* 2011; Sánchez-Sevilla *et al.* 2015; Hancock *et al.* 2018) but none have painted a holistic picture of the complicated wild ancestry and dynamic forces that shaped genetic diversity in *F. × ananassa*.

**Table 1.** Number of Primary, Secondary, and Tertiary Gene Pool Founders in the Global Genealogy of Cultivated Strawberry.

| Species | Ploidy | Giant | Halo | Complete |
| --- | --- | --- | --- | --- |
| **Primary Gene Pool** |  |  |  |  |
| *F. chiloensis* | 2n = 8x = 56 | 79 | 33 | 112 |
| *F. virginiana* | 2n = 8x = 56 | 41 | 24 | 65 |
| *F. × ananassa* | 2n = 8x = 56 | 656 | 515 | 1,171 |
| Unknown Octoploid *Fragaria* | 2n = 8x = 56 | 9 | 1 | 10 |
| **Primary Gene Pool Total** |  | **785** | **573** | **1,358** |
| **Secondary Gene Pool** |  |  |  |  |
| *F. iinumae* | 2n = 2x = 14 | 1 | 2 | 3 |
| *F. nilgerrensis* | 2n = 2x = 14 | 2 | 0 | 2 |
| *F. nipponica* | 2n = 2x = 14 | 0 | 2 | 2 |
| *F. nubicola* | 2n = 2x = 14 | 2 | 0 | 2 |
| *F. orientalis* | 2n = 2x = 14 | 3 | 1 | 4 |
| *F. viridis* | 2n = 2x = 14 | 4 | 2 | 6 |
| *F. vesca* | 2n = 2x = 14 | 20 | 24 | 44 |
| *F. moschata* | 2n = 6x = 42 | 6 | 0 | 6 |
| *F. × vescana* | 2n = 10x = 70 | 1 | 0 | 1 |
| **Secondary Gene Pool Total** |  | **39** | **31** | **70** |
| **Tertiary Gene Pool** |  |  |  |  |
| *P. glandulosa* | 2n = 2x = 14 | 3 | 0 | 3 |
| *P. anserina* | 2n = 4x = 28 | 1 | 0 | 1 |
| *P. palustris* | 2n = 6x = 42 | 1 | 4 | 5 |
| Unknown *Potentilla* | NA | 0 | 1 | 1 |
| **Tertiary Gene Pool Total** |  | **5** | **5** | **10** |

We identified 1,438 founders in the genealogy of cultivated strawberry (Fig. 1; Table 1; Files S1, S4-S5). Here and elsewhere, ‘founders’ are individuals with unknown parents, whereas ‘ancestors’ are ascendants that may or may not be founders (Lacy 1989, 1995). The terminal nodes in the pedigree networks are either founders or the youngest descendants in a pedigree (Figs. 1-2). Of the 1,438 founders, 267 were wild species and 1,171 were *F. × ananassa* individuals (Fig. 1; Table 1). Because the *F. × ananassa* founders are either interspecific hybrids or descendants of interspecific hybrids, the number of wild species founders could exceed 268. One of the challenges we had with estimating the number of wild species founders was the anonymity of ecotypes that were used as parents before breeders began carefully documenting pedigrees (File S1). We could not rule out that some of the anonymous wild species founders in the pedigree records might have been clones of the same individuals, which means that the estimated number of wild species founders reported here could be inflated.

As interspecific hybridization with wild founders became less important and intraspecific (*F. × ananassa*) hybridization became more important in strawberry breeding, the proportional genetic contribution of wild founders to the gene pool of cultivated strawberry decreased (Fig. 5; Files S4-S5). This seems paradoxical because 100% of the alleles found in *F. × ananassa* were inherited from wild founders, but increasingly flowed through *F. × ananassa* descendants over time—wild octoploids numerically only constituted 14% of the founders we identified (Table 1). Several trends emerged from our analyses of genetic relationships and founder contributions. First, inbreeding has steadily increased over time as a consequence of population bottlenecks and directional selection (Fig. 5B). Second, the California population was significantly more inbred than the cosmopolitan population (Fig. 5B). These results were consistent with the findings of Hardigan *et al.* (2020b) from genome-wide analyses of DNA variants and population structure. They found selective sweeps on several chromosomes in the California population, which was shown to be unique and bottlenecked. Finally, the relative number of founder equivalents (Lacy 1989, 1995) has decreased over time, consistent with the increase in inbreeding over time (Fig. 5A-B).

**Figure 5.**
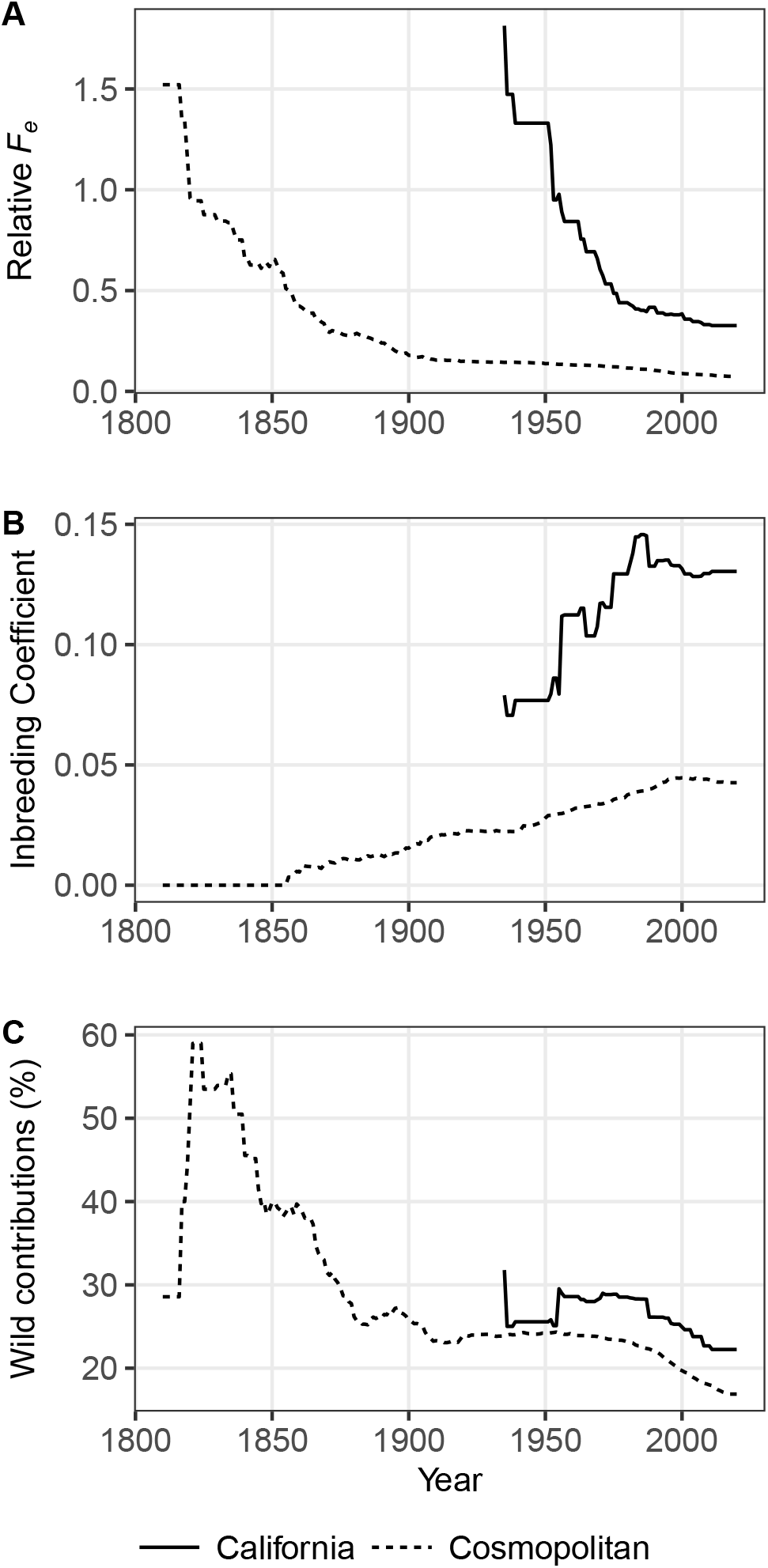
Relative Founder Equivalents, Inbreeding Coefficients, and Wild Founder Genetic Contributions Over Time. (A) Relative founder equivalent (*F*_*e*_/*n*) estimates for California and cosmopolitan cultivars over time, where *F*_*e*_ = founder equivalents and *n* = number of founders. The California population included 69 cultivars developed at the University of California, Davis (UCD) since the inception of the breeding program in 1924. The birth year (year of origin) was known for all of the UCD cultivars. The cosmopolitan population included 2,140 cultivars with known birth years. (B) Wright’s coefficient of inbreeding (*F*) for individuals in the California and cosmopolitan populations over time. *F* was estimated from the relationship matrix (*A*). (C) Estimates of the genetic contributions of wild species founders to allelic diversity in the California and cosmopolitan populations.

### Primary, Secondary, and Tertiary Gene Pool Founders of Cultivated Strawberry

Predictably, the wild species founders of *F. × ananassa* were dominated by *F. chiloensis* (*n* = 112) and *F. virginiana* (*n* = 65) (Table 1). Seven of eight subspecies of *F. chiloensis* and *F. virginiana* (Staudt 1988; Hummer *et al.* 2011) were identified in pedigree records: *F. chiloensis* subsp. *chiloensis*, *F. chiloensis* subsp. *lucida*, *F. chiloensis* subsp. *pacifica*, and *F. chiloensis* subsp. *sandwicensis*, *F. virginiana* subsp. *virginiana*, *F. virginiana* subsp. *glauca*, and *F. virginiana subsp. platypetala* (Bringhurst 1918-2016; https://www.ars.usda.gov/; Fig. 1; Table 1; File S1). Primary gene pool individuals (187 wild octoploid ecotypes and 1,171 hybrid *F. × ananassa* individuals) constituted 95% of the founders and were estimated to have contributed ≥ 99% of the allelic diversity to global, California, and cosmopolitan *F. × ananassa* populations (Fig. 6; Table 1; Files S4-S5). Even though wild species from the secondary (*n* = 70) and tertiary (*n* = 10) gene pools of *F. × ananassa* constituted 6% of the founders and 30% of the wild species founders identified in pedigree records, they were estimated to have contributed < 0.1% of the allelic diversity in the global *F. × ananassa* population (Table 1; Files S4-S5).

**Figure 6.**
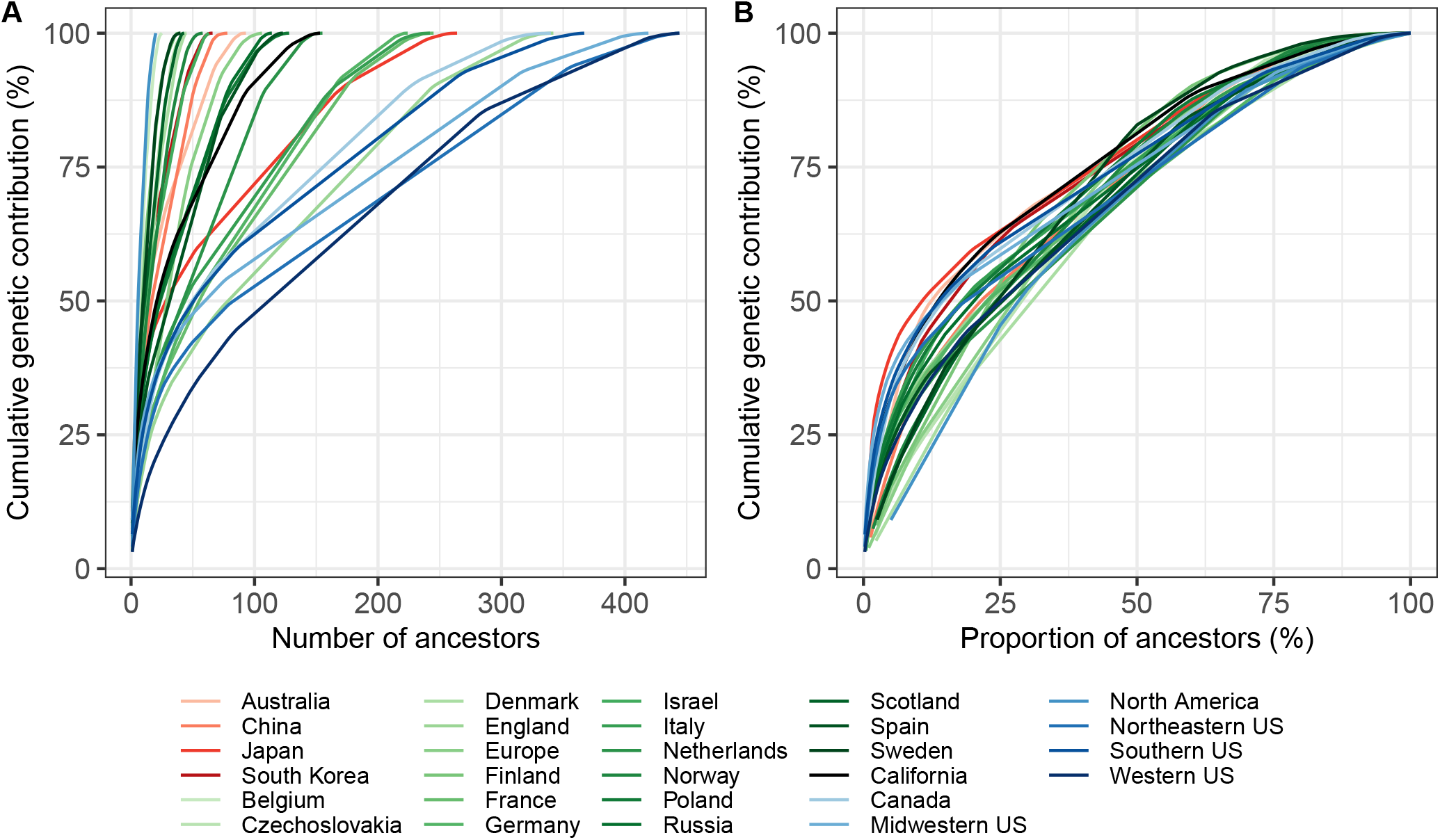
Genetic Contributions of Ancestors to Cultivars. (A) The genetic contributions of ancestors to the allelic diversity among *k* cultivars within a focal population were estimated from the mean coancestry between the *i*th ancestor and the *k* cultivars within the focal population. The genetic contributions of the ancestors were ordered from largest to smallest to calculate the cumulative genetic contributions of ancestors to cultivars in a focal population. (B) The proportion of ancestors needed to account for *p*% of the allelic diversity among cultivars within a focal population was estimated by dividing the cumulative genetic contribution by *k*.

The secondary and tertiary gene pool founders were primarily parents of orphans or other isolated individuals in short dead-end pedigrees that have not materially contributed allelic diversity to the primary gene pool. These included decaploid (2*n* = 10*x* = 70) *F. × vescana* and pentaploid (2*n* = 5*x* = 35) *F. × bringhurstii* individuals (Bringhurst and Senanayake 1966; Bauer 1994; Sangiacomo and Sullivan 1994; Hummer *et al.* 2011). Although frequently cited as important genetic resources for strawberry breeding (Darrow 1966; Hummer 2008; Gaston *et al.* 2020), the secondary and tertiary gene pools of cultivated straw-berry have had limited utility because of the range of biological challenges one encounters when attempting to introgress alleles from exotic sources through interspecific and intergeneric hybrids, e.g., reproductive and recombination barriers, ploidy differences, meiotic abnormalities, and hybrid sterility (Bringhurst and Senanayake 1966; Bringhurst and Gill 1970; Harlan and de Wet 1971; Evans 1977; Bauer 1994; Sangiacomo and Sullivan 1994).

The secondary and tertiary gene pools are hardly needed to drive genetic gains or solve problems in strawberry breeding. Hardigan *et al.* (2020b) showed that genetic diversity is massive in the primary gene pool and has not been eroded by domestication and breeding on a global scale, even though it has been significantly reduced and restructured in certain populations, e.g., the California population. The profound changes and re-structuring in the California population over time, as previously noted, were clearly evident in the sociograms and PCAs of the pedigree-genomic relationship matrices (Figs. 1-2). Because the California population has been the source of numerous historically and commercially important cultivars, we hypothesize that intense selection and population bottlenecks have purged a high frequency of unfavorable alleles compared to many other populations, thereby yielding an elite population with lower genetic diversity than the highly admixed cosmopolitan population (Figs. 1-2; Hardigan *et al.* 2020b).

### Prominent and Historically Important Ancestors of Cultivated Strawberry

We used coancestry, betweenness centrality (*B*), and out-degree (*d*_*o*_) statistics to estimate the genetic contribution (*GC*) of founders and non-founders to genetic variation within a popu-lation and identify the most prominent and important ancestors in the genealogy of cultivated strawberry (Freeman 1977; Scott 1988; Lacy 1989, 1995; Fig. 6; Table 2; Files S4-S5). The estimation of *GC* from the coancestry matrix (*A*) differed between founders and ancestors (founders and non-founders). For founders, *GC* was estimated by the mean coancestry or kinship (*MK*) between each founder and cultivars within a focal population (Files S4-S5). For ancestors, *GC* was iteratively estimated by *MK* between each ancestor and cultivars within a focal population, starting with the ancestor with the largest *MK* estimated from *A*, deleting that ancestor, re-estimating the coancestry matrix (*A^*^*), selecting the ancestor with the largest *MK* estimated from the pruned coancestry matrix (*A^*^*), deleting that ancestor, re-estimating the coancestry matrix, and repeating until every ancestor had been dropped. We compiled *GC*, *B*, and *d*_*o*_ estimates for every founder and non-founder in the pedigree database (Files S4-S5).

**Table 2.** The Twenty-Most Prominent and Historically Important Ancestors of Cultivars.

| California |  |  |  | Cosmopolitan |  |  |  |
| --- | --- | --- | --- | --- | --- | --- | --- |
| Ancestor | GC (%) | B | d<sub>o</sub> | Ancestor | GC (%) | B | d<sub>o</sub> |
| Tufts | 12.2 | 52,013.9 | 80 | Howard 17 | 4.4 | 47,942.5 | 99 |
| Lassen | 7.1 | 56,157.0 | 42 | Fairfax | 1.9 | 13,090.4 | 91 |
| Cal 177.21 | 6.4 | 36,728.6 | 49 | Hovey | 1.8 | 12,390.6 | 19 |
| Douglas | 5.7 | 72,781.8 | 32 | Tufts | 1.4 | 16,579.3 | 12 |
| 71C098P605 | 3.6 | 16,434.8 | 13 | Crescent | 1.3 | 16,803.7 | 59 |
| Nich Ohmer | 3.0 | 2,977.0 | 124 | Aberdeen | 1.2 | 7,908.6 | 35 |
| Camino Real | 2.6 | 17,797.1 | 23 | Sharpless | 1.2 | 11,727.0 | 51 |
| Howard 17 | 2.5 | 52,231.1 | 16 | Blakemore | 1.2 | 13,265.9 | 49 |
| Sequoia | 2.4 | 40,254.5 | 38 | Wilson | 1.0 | 4,012.6 | 51 |
| Diamante | 2.3 | 31,032.9 | 27 | Royal Sovereign | 0.9 | 19,373.0 | 23 |
| Irvine | 2.0 | 11,938.8 | 12 | Harunoka | 0.9 | 6,193.6 | 24 |
| Palomar | 1.9 | 27,644.3 | 22 | Douglas | 0.8 | 22,433.6 | 23 |
| Albion | 1.8 | 22,016.6 | 11 | Gorella | 0.7 | 12,053.2 | 41 |
| 42C008P016 | 1.8 | 12,687.4 | 26 | Hoffman | 0.7 | 5,738.0 | 17 |
| Parker | 1.5 | 2,924.8 | 10 | Marshall | 0.7 | 0.0 | 58 |
| 65C065P601 | 1.5 | 19,867.1 | 13 | Holiday | 0.6 | 6,157.4 | 39 |
| Seascape | 1.5 | 8,637.0 | 12 | Senga Sengana | 0.6 | 3,258.0 | 58 |
| San Andreas | 1.3 | 35,857.9 | 22 | Bubach | 0.6 | 0.0 | 56 |
| Aiko | 1.2 | 8,141.0 | 5 | Reiko | 0.6 | 2,766.0 | 19 |
| Oso Grande | 1.1 | 48,118.7 | 20 | Cumberland Triumph | 0.5 | 10,544.7 | 12 |

| California |  |  |  | Cosmopolitan |  |  |  |
| --- | --- | --- | --- | --- | --- | --- | --- |
| Ancestor | GC (%) | *B* | *d<sub>o</sub>* | Ancestor | GC (%) | *B* | *d<sub>o</sub>* |
| Tufts | 12.2 | 52,013.9 | 80 | Howard 17 | 4.4 | 47,942.5 | 99 |
| Lassen | 7.1 | 56,157.0 | 42 | Fairfax | 1.9 | 13,090.4 | 91 |
| Cal 177.21 | 6.4 | 36,728.6 | 49 | Hovey | 1.8 | 12,390.6 | 19 |
| Douglas | 5.7 | 72,781.8 | 32 | Tufts | 1.4 | 16,579.3 | 12 |
| 71C098P605 | 3.6 | 16,434.8 | 13 | Crescent | 1.3 | 16,803.7 | 59 |
| Nich Ohmer | 3.0 | 2,977.0 | 124 | Aberdeen | 1.2 | 7,908.6 | 35 |
| Camino Real | 2.6 | 17,797.1 | 23 | Sharpless | 1.2 | 11,727.0 | 51 |
| Howard 17 | 2.5 | 52,231.1 | 16 | Blakemore | 1.2 | 13,265.9 | 49 |
| Sequoia | 2.4 | 40,254.5 | 38 | Wilson | 1.0 | 4,012.6 | 51 |
| Diamante | 2.3 | 31,032.9 | 27 | Royal Sovereign | 0.9 | 19,373.0 | 23 |
| Irvine | 2.0 | 11,938.8 | 12 | Harunoka | 0.9 | 6,193.6 | 24 |
| Palomar | 1.9 | 27,644.3 | 22 | Douglas | 0.8 | 22,433.6 | 23 |
| Albion | 1.8 | 22,016.6 | 11 | Gorella | 0.7 | 12,053.2 | 41 |
| 42C008P016 | 1.8 | 12,687.4 | 26 | Hoffman | 0.7 | 5,738.0 | 17 |
| Parker | 1.5 | 2,924.8 | 10 | Marshall | 0.7 | 0.0 | 58 |
| 65C065P601 | 1.5 | 19,867.1 | 13 | Holiday | 0.6 | 6,157.4 | 39 |
| Seascape | 1.5 | 8,637.0 | 12 | Senga Sengana | 0.6 | 3,258.0 | 58 |
| San Andreas | 1.3 | 35,857.9 | 22 | Bubach | 0.6 | 0.0 | 56 |
| Aiko | 1.2 | 8,141.0 | 5 | Reiko | 0.6 | 2,766.0 | 19 |
| Oso Grande | 1.1 | 48,118.7 | 20 | Cumberland Triumph | 0.5 | 10,544.7 | 12 |

We identified four *F. chiloensis*, five *F. virginiana*, and 40 *F. × ananassa* founders in the genealogy of the California population (File S4). Cumulative *GC* estimates for the California population were 1.8% for *F. chiloensis*, 12.7% for *F. virginiana*, and 85.5% for *F. × ananassa* founders. Four of the nine wild octoploid founders of the California population were founders of the historic Ettersburg population that supplied genetic diversity for private and public sector breeding programs in California (Clausen 1915; Wilhelm and Sagen 1974; Bringhurst *et al.* 1990; Sjulin 2006). The wild octoploid founders with the largest genetic contributions were three *F. virginiana* ecotypes: ‘New Jersey Scarlet’ (8.3%), ‘Hudson Bay’ (2.7%), and ‘Wasatch’ (1.3%) (Table S1). Wasatch is the *F. virginiana* subsp. *glauca* donor of the *PERPETUAL FLOW-ERING* mutation that Bringhurst *et al.* (1980) transferred into *F. × ananassa* (Bringhurst *et al.* 1989). The Wasatch ecotype appears in the genetic background of every day-neutral cultivar developed at the University of California, Davis. Similarly, we identified 26 *F. chiloensis*, 24 *F. virginiana*, and 490 *F. × ananassa* founders in the genealogy of the cosmopolitan population (File S5). Cumulative *GC* estimates for the cosmopolitan population were 4.6% for *F. chiloensis*, 14.1% for *F. virginiana*, 79.9% for *F. × ananassa*, and 1.4% for other founders. Similar to what we found for the California population, the wild octoploid founders with the largest genetic contributions were ‘New Jersey Scarlet’ (8.3%) and ‘Hudson Bay’ (3.5%) (Fletcher 1917; Darrow 1937). The next largest genetic contribution was made by FC_071 (1.9%), an *F. chiloensis* ecotype of unknown origin found in the pedigrees of Madame Moutot, Sharpless, Royal Sovereign, and other influential early cultivars (Table S1; Figure 4).

A significant fraction of the alleles found in *F. × ananassa* populations have flowed through a comparatively small number of common ancestors, each of which have contributed unequally to standing genetic variation (Fig. 6; Table 2; Files S4-S5). The most important ancestors are described as ‘stars’ in the lexicon of social network analysis, and are either locally or globally central (Moreno 1953; Scott 1988; Wasserman and Faust 1994). Globally central individuals reside in the upper right quadrant of the *B* × *d*_*o*_ distribution 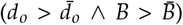, where 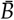 is the mean of *B* and 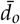 is the mean of *d*_*o*_ –8.7-8.9% of the ancestors were classified as globally central (Fig. 7; Moreno 1953; Scott 1988; Wasserman and Faust 1994). Locally central individuals reside in the upper left quadrant of the *B* × *d*_*o*_ distribution 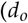 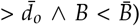 —11.8-12.1% of the ancestors were classified as locally central (Fig. 7; Moreno 1953; Scott 1988; Wasserman and Faust 1994). ‘Tufts’, ‘Lassen’, ‘Nich Ohmer’, ‘Howard 17’, and ‘Fairfax’ were among the biggest stars, along with several other iconic, mostly heirloom cultivars, and all were either locally or globally central (Table 2). Stars are ‘gatekeepers’ that have numerous descendants (the largest *d*_*o*_ estimates), transmitted a disproportionate fraction of the alleles found in a population (have the largest *GC* estimates), have the largest number of inter-connections (largest *B* estimates) in the pedigree, and are visible in sociograms as nodes with radiating pinwheel-shaped patterns of lines (Fig 2; Table 2; Files S4-S5). Several of the latter are visible in the sociograms we developed for the California and cosmopolitan populations. Stars have the largest nodes (*B* estimates) in the sociograms (Fig 2).

**Figure 7.**
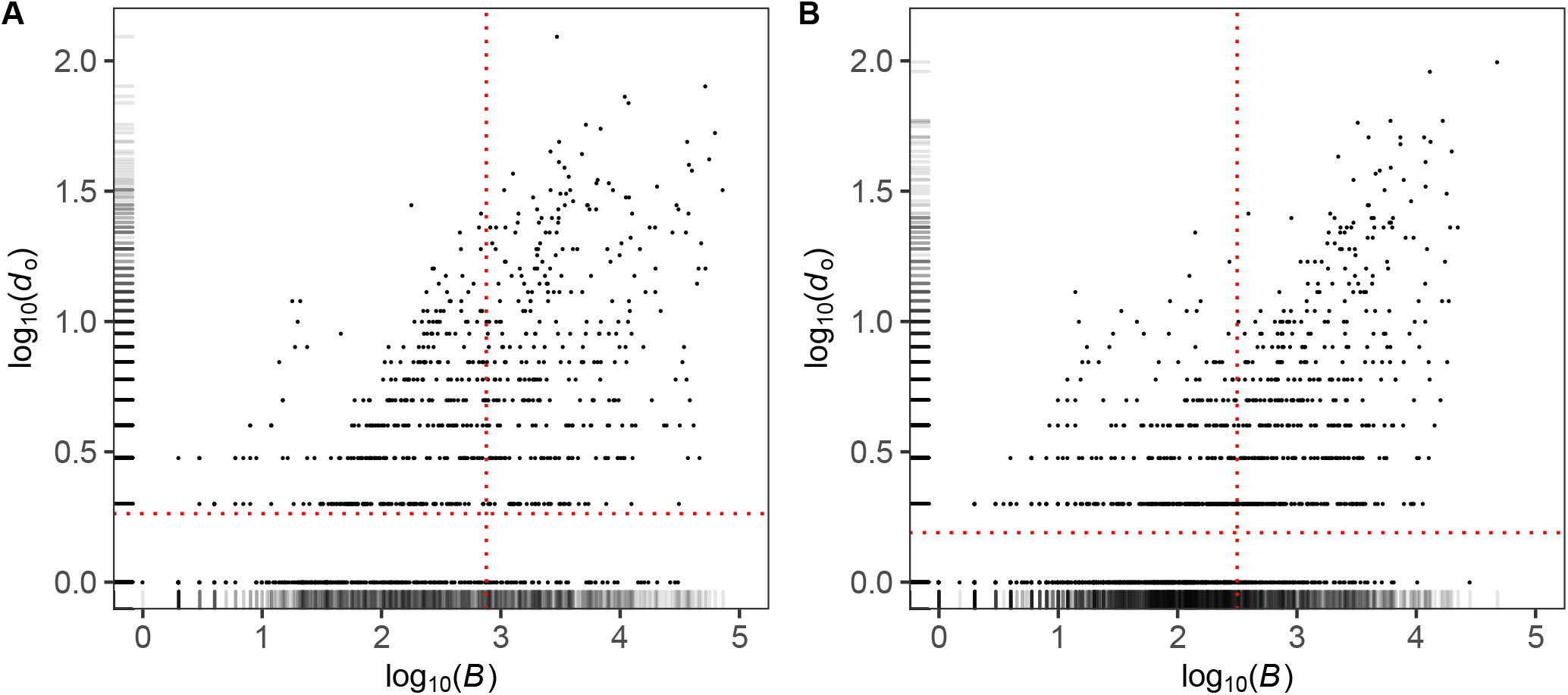
Structural Roles and Betweenness Centrality (*B*) and Out-Degree (*d_o_*) Statistics for Individuals in Cultivated Strawberry Sociograms. (A) *B* and *d_o_* estimates for individuals in the California population. (B) *B* and *d*_*o*_ estimates for individuals in the cos-mopolitan population. (A) and (B) The red dashed lines delineate globally central (upper right; 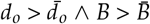), locally central (upper left; 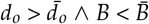), broker (lower right; 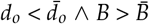), and marginal (lower left; 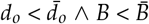) quadrants, where 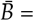 the mean of *B* estimates and 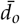 = the mean of *d*_*o*_ estimates. 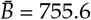 and 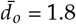 for the California population, whereas 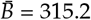 and 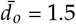 for the cosmopolitan population. *B* and *d*_*o*_ estimate densities are plotted along the x- and y-axes.

We estimated and compiled *GC* statistics for every ancestor in the California and cosmopolitan populations (Files S4-S5). The twenty-most prominent and historically important ancestors of the California and cosmopolitan populations are shown in Table 2. They include several iconic and well known heirloom and modern cultivars, e.g., ‘Tioga’, ‘Douglas’, and ‘Royal Sovereign’ (Fletcher 1917; Darrow 1937, 1966; Wilhelm and Sagen 1974; Sjulin and Dale 1987; Bringhurst *et al.* 1990), in addition to ‘un-released’ germplasm accessions preserved in the UCD Straw-berry Germplasm Collection, e.g., 65C065P601 (aka 65.65-1). The latter is the oldest living descendant of the aforementioned *F. virginiana* subsp. *glauca* ‘Wasatch’ ecotype collected by Royce S. Bringhurst from the Wasatch Mountains, Little Cottonwood Canyon, Utah (Bringhurst *et al.* 1980, 1989; Ahmadi *et al.* 1990). The ‘Wasatch’ ecotype is a founder of every day-neutral cultivar in the California population and many day-neutral cultivars in the cosmopolitan population with alleles flowing through 65C065P601 and the UCD cultivar ‘Selva’ (Bringhurst *et al.* 1989; Files S4-S5).

*GC* statistics were ordered from largest to smallest (*GC*_1_ ≥ *GC*_2_ ≥… ≥ *GC*_*n*_) and progressively summed to calculate the cumulative genetic contributions of ancestors and the number of ancestors needed to explain *p*% of the genetic variation (*n*_*p*_) in a focal population, where *p* ranges from 0 to 100% (Fig. 6). The parameter *n*_100_ estimates the number of ancestors needed to account for 100% of the genetic variation among *k* cultivars in a focal population (each focal population was comprised of cultivars, ascendants, and descendants). *n*_100_ estimates were 153 for the California population and 3,240 for the cosmopolitan population. The latter number was significantly larger than the number for the California population because the cosmopolitan population includes pedigrees for 2,499 cultivars developed worldwide, whereas the California population includes pedigrees for 69 UCD cultivars only (File S1). Within European countries, *n*_100_ ranged from 25 for Belgium to 342 for England (Fig. 6A). Within the US, *n*_100_ ranged from a minimum of 367 for the southern region to a maximum of 444 for western and northeastern regions.

Predictably, *n*_*p*_ increased at a decreasing rate as the number of *GC*-ranked ancestors increased (Fig. 6). Cumulative *GC* estimates increased as non-linear diminishing-return functions of the number of ancestors (Table 2; Files S4-S5). The slopes were initially steep because a fairly small number of ancestors accounted for a large fraction of the genetic variation within a particular focal population. Across continents, regions, and countries, eight to 112 ancestors accounted for 50% of the allelic variation within focal populations (Fig. 6; Table 2). The differences in *n*_*p*_ estimates were partly a function of the number of cultivars (*k*) within each focal population. When *n*_*p*_ was expressed as a function of *k*, we found that the proportion of ancestors needed to explained *p*% of the allelic variation in a focal population was strikingly similar across continents, regions, and countries, e.g., the Western US population, which had the largest *n*_100_ estimate (Fig. 6A), fell squarely in the middle when expressed as a function of *k* (Fig. 6B).

### Breeding Speed in Cultivated Strawberry

Social network analyses of the pedigree networks shed light on the speed of breeding and changes in the speed of breeding over the last 200 years in strawberry (Figs. 8-9). We retraced the ancestry of every cultivar through nodes and edges in the sociograms (Figs. 1-2). The year of origin was known for 71% of the individuals. These edges yielded robust estimates of the mean selection cycle length in years (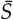 = mean number of years/generation). 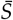 was calculated from thousands of directed acyclic graphs, which are unidirectional paths traced from culti-vars back through descendants to founders (Thulasiraman and Swamy 1992). Collectively, cultivars in the California population (*n* = 69) visited 27,058 parent-offspring edges, whereas culti-vars in the cosmopolitan population (*n* = 1,982) visited 155,487 parent-offspring edges. The selection cycle length means (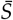) and distributions over the last 200 years were strikingly similar across continents, regions, and countries—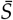 was 16.9 years/generation for the California population and 16.0 years/generation for the cosmopolitan population (Fig. 8). These extraordinarily long selection cycle lengths are more typical of a long-lived woody perennial than a fast cycling annual (van Nocker and Gardiner 2014; Jighly *et al.* 2019); however, the speed of breeding has steadily increased over time (Fig. 8). By 2000, 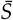 had decreased to six years/generation in the California population and 10 years/generation in the cosmopolitan population (Fig. 9).

**Figure 8.**
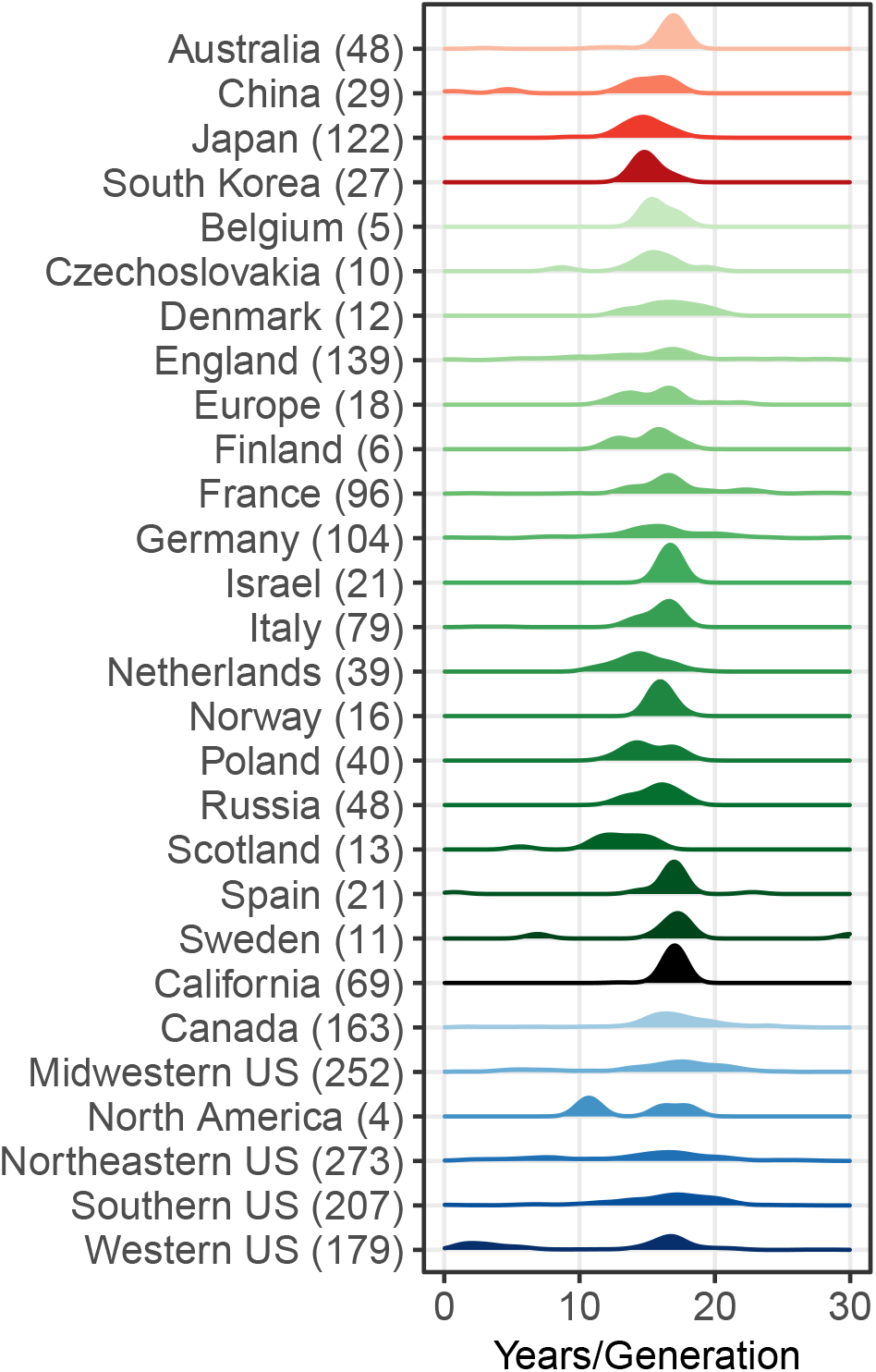
Selection Cycle Length Distributions by Geography. Selection cycle length means (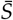 = mean number of years/generation) were estimated for *k* cultivars within continent-, region-, and country-specific focal populations of cultivated strawberry (*k* is shown in parentheses for each geographic group). 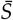 was estimated from edge lengths (years/edge) for all possible paths (directed graphs with alleles flowing from parents to offspring but not *vice versa*) in pedigrees connecting cultivars to founders, where the length of an edge = the birth year difference between parent and offspring. 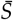 probability densities are shown for cultivars developed in different countries, regions, or continents. Only estimates in the zero to 30 year/generation range are shown because estimates exceeding 30 years/generation were extremely rare.

**Figure 9.**
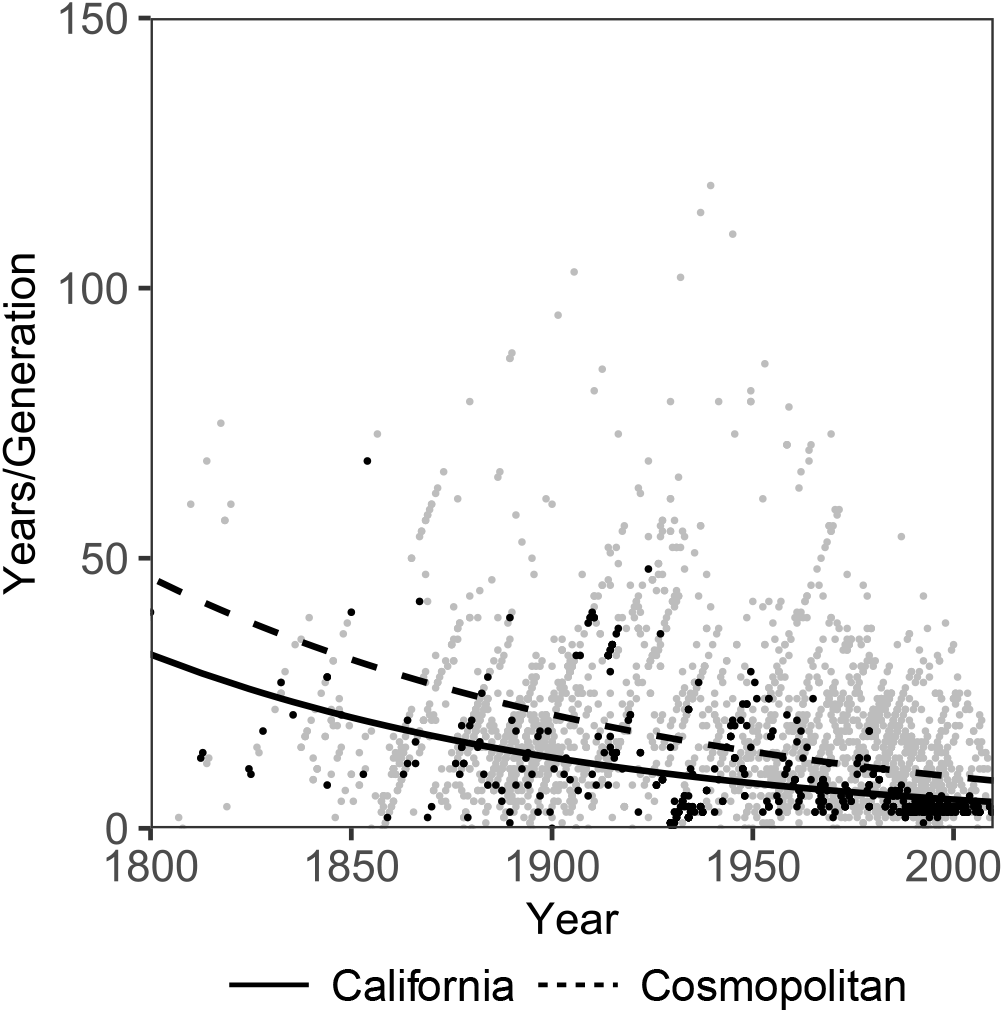
Breeding Speed Over Time. Selection cycle lengths (*S* = years/generation) were estimated for 3,693 independent parent-offspring edges in the pedigree networks for the Califor-nia and cosmopolitan populations. *S* estimates were limited to parents and offspring with known birth years. Selection cycle lengths are plotted against the midpoint (*m*) between parent and offspring birth years for California (black points) and cosmopolitan (gray points) populations. The plotted lines are exponential decay functions fitted by non-linear regression of *S* on *m*. The function for the California population was *y* = 35.06 · *e*^−0.0090·(*x*−1790.5)^ (Nagelkerke pseudo-*R*^2^ = 0.25; *p* < 0.001). The function for the cosmopolitan population was *y* = 76.69 · *e*^−0.0079·(*x*−1736.5)^ (Nagelkerke pseudo-*R*^2^ = 0.08; *p* < 0.001).

The genealogy does not account for lineages underlying what must have been millions of hybrid progeny screened in breeding programs worldwide, e.g., Johnson (1990) alone reported screening 600,000 progeny over 34 years (1956-1990) at Driscoll’s (Watsonville, California). Cultivars are nevertheless an accurate barometer of global breeding activity and the only outward facing barometer of progress in strawberry breeding. When translated across the last 200 years of breeding, our selection cycle length estimates imply that the 2,656 cultivars in the genealogy of cultivated strawberry have emerged from the mathematical equivalent of only 12.9 cycles of selection (200 years 15.5 years per generation). Even though offspring from 250 years of crosses have undoubtedly been screened worldwide since 1770, 15.5 years have elapsed on average between parents and offspring throughout the history of strawberry breeding (Fig. 8-9). Because genetic gains are affected by selection cycle lengths, and faster generation times normally translate into greater genetic gains and an increase in the number of recombination events per unit of time (Bernardo 2002; Ceccarelli 2015; Bernardo 2017; Jighly *et al.* 2019; Bernardo 2020), our analyses suggest that genetic gains can be further increased in strawberry by shortening selection cycle lengths. Genome-informed breeding, speed breeding, and other technical innovations are geared towards that goal and have the potential to shorten selection cycle lengths and increase genetic gains (van Nocker and Gardiner 2014; Whitaker *et al.* 2020).

## Acknowledgements

This research was supported by grants to SJK from the United Stated Department of Agriculture (http://dx.doi.org/10.13039/100000199) National Institute of Food and Agriculture (NIFA) Specialty Crops Research Initiative (#2017-51181-26833), California Strawberry Commission (http://dx.doi.org/10.13039/100006760), and the University of California, Davis (http://dx.doi.org/10.13039/100007707) and to SLD from the National Science Foundation Division Of Integrative Organismal Systems (#1444478). The USDA grant supported the dissertation research of DDP and MJF. The postdoctoral research of CH was supported by the NSF grant. We are grateful to Clint Pumphrey, the manuscript curator of the special collections and archives of the Merill-Cazier Library at Utah State University (Logan, Utah). Clint assisted the first author with acquiring and researching the laboratory notebooks and other records of Royce S. Bringhurst (1918-2005), a former faculty member and strawberry breeder at the University of California, Davis (1953-1989). The documents and photos associated with the collection yielded extensive pedigree records that were crucial for reconstructing the genealogy of the UCD Strawberry Breeding Program. We are equally grateful to Phillip Stewart, a strawberry breeder at Driscoll’s (Watsonville, California), for sharing copies of the University of California, Berkeley (UCB) pedigree records of Harold E. Thomas (1900-1986), a former faculty member and strawberry breeder at UCB from 1927 to 1945. Those pedigree records greatly increased the completeness and depth of the database for the early years of the University of California Strawberry Breeding Program. The authors thank Thomas Sjulin, a former strawberry breeder at Driscoll’s (Watsonville, California), for sharing the public pedigree records he assembled over his career. Those nucleated the pedigree database we developed and were a catalyst for our study. SJK and GSC thank Robert Kerner (Information Technology Manager, Department of Plant Sciences, UCD) for the computer forensic analyses that recovered several hundred pedigree records for UCD individuals from an obsolete electronic database, thus preventing the loss of those records for perpetuity. They were critical for integrating the UCD genealogy with the global genealogy for cultivated strawberry. SJK especially thanks Rachel Krevans, Matthew Chivvus, Jake Ewert, and Wesley Overson (lawyers at Morrison-Forester, San Francisco, California) for their integrity, friendship, and steadfast support.

